# MONTE enables serial immunopeptidome, ubiquitylome, proteome, phosphoproteome, acetylome analyses of sample-limited tissues

**DOI:** 10.1101/2021.06.22.449417

**Authors:** Jennifer G. Abelin, Erik J. Bergstrom, Hannah B. Taylor, Keith D. Rivera, Susan Klaeger, Charles Xu, C. Jackson White, Meagan E. Olive, Myranda Maynard, M. Harry Kane, Suzanna Rachimi, D.R. Mani, Michael A. Gillette, Karl R. Clauser, Namrata D. Udeshi, Steven A. Carr

## Abstract

Serial multiomic analyses of proteome, phosphoproteome and acetylome provides functional insights into disease pathology and drug effects while conserving precious human material. To date, ubiquitylome and HLA peptidome analyses have required separate samples for parallel processing each using distinct protocols. Here we present MONTE, a highly-sensitive **m**ulti-**o**mic **n**ative **t**issue **e**nrichment workflow that enables serial, deepscale analysis of HLA-I and HLA-II immunopeptidome, ubiquitylome, proteome, phosphoproteome and acetylome from the same tissue samples. We demonstrate the capabilities of MONTE in a proof-of-concept study of primary patient lung adenocarcinoma(LUAD) tumors. Depth of coverage and quantitative precision at each of the ‘omes is not compromised by serialization, and the addition of HLA immunopeptidomics enables identification of putative immunotherapeutic targets such as cancer/testis antigens and neoantigens. MONTE can provide insights into disease-specific changes in antigen presentation, protein expression, protein degradation, cell signaling, cross-talk and epigenetic pathways involved in disease pathology and treatment.

## Introduction

Mass spectrometry-based proteomics is a proven technology for characterization of cell surface immunopeptidomes along with intracellular cellular proteins and their post-translational modifications (PTMs)^1–10^. Technological advances to measure each of these ‘omes to deepen our understanding of tumor biology, infectious diseases, autoimmunity, and other diseases have been made. However, patient tissues are often limited in quantity. One of the main ways to overcome this limitation has been to serialize sample processing such that the flow-through of one enrichment step (e.g., phosphopeptide enrichment) is used as the input for the next enrichment step (e.g., acetylpeptide enrichment). Current serial enrichment workflows for measuring ‘omes at high multiplex leverage isobaric reagents such as TMT^11–15^ or iTRAQ^16,17^. However, to date it has not been possible to include either immunopeptidome or ubiquitylome (i.e. anti-K-ε-GG antibody enrichment) in serial. This is because conventional immunopeptidomics and ubiquitylomics protocols require separate sample aliquots as they cannot be enriched after chemical labeling because the immunopeptidome enrichment is done prior to sample digestion and enrichment of ubiquitylated peptides occurs prior to TMT labeling^3,10,18,19^. These limitations impede the identification of cancer driver signatures and the detection of signaling network adaptations, changes in antigen presentation, molecular complex formation and protein localization by concordant readout of the immunopeptidome, proteome and PTM-omes(ubiquitylome, phosphoproteome, acetylome).

There are two main complications that preclude adding immunopeptidome analysis in a serial processing strategy. The first is that much more sample input has typically been used for immunopeptidomics than for proteomic and PTM-omics in order to enable detection of low abundant, clinically relevant antigens such as neoantigens^2,6^. For immunopeptidomics, a separate aliquot of tissue, usually 500 to 1000 milligrams of wet weight tissue is needed compared to 25-50 milligrams for serial, multiplexed proteomics, phospho- and acetylpeptidomics^11–14,17^. Moreover, sample preparation for immunopeptidomics is distinct from that used for conventional proteomics. Immunopurification(IP) of HLA molecules requires the use of native lysis buffer containing mild detergent to maintain protein conformations and solubilize membrane bound HLA proteins. In contrast, current serial proteome and PTM-ome enrichment protocols denature proteins using urea or SDS prior to tryptic digestion preventing upstream HLA peptide complex enrichment.

Historically, the ubiquitylome has also been difficult to analyze in high-multiplex for primary tissue samples due to the inability of the antibody used for enrichment of di-glycyl remnant (K-ε-GG) of formerly ubiquitylated peptides to recognize these peptides once the newly generated N-terminal glycine is labeled with TMT or iTRAQ reagents^10,18^. To address this issue, we recently developed^3^ and automated^20^ UbiFast, a highly-sensitive, rapid and multiplexed protocol for quantifying >10,000 ubiquitylation sites from cells or tissue in a TMT10plex^3^. UbiFast^20^ labels K-ε-GG peptides with TMT reagents while the modified peptides are still bound to the anti-K-ε-GG antibody. The key to the UbiFast approach is the on-antibody labeling step that allows TMT labeling of peptide N-terminal amine groups and the ε-amine groups of lysine residues, but not the primary amine of the di-glycyl remnant which is protected from labeling by the antibody. Because the UbiFast method requires enrichment of K-ε-GG peptides prior to TMT labeling and sample mixing, the method has only been used to enrich ubiquitylated peptides from a separate sample aliquot in parallel to samples processed for whole proteome and other PTM-omes[Citation error].

To overcome the challenges of serializing deep-scale immunopeptidome and ubiquitylome with proteome, phosphoproteome and acetylome profiling from a single tissue sample of limited quantity, we have developed an integrated proteomics workflow that we term MONTE (Multi-Omic Native Tissue Enrichment). MONTE integrates our published methods for isolation of immunoprecipitated HLA peptide complexes from as few as 50 million cells and 0.2g of clinical specimens^7,8,21,22^, in combination with recent improvements in MS instrumentation, off-line fractionation, and separation in the gas phase^23^ to increase immunopeptidome yield. The flow-through of the HLA immunopeptidome purification contains the intact cellular proteome that we make compatible with the current multiplexed, serialized multi-proteomics-workflow by use of SDS based lysis and tryptic digestion post HLA enrichment^24^. The resulting digest is then processed and analyzed by the UbiFast workflow for multiplexed ubiquitylation profiling using anti-K-ε-GG antibodies and on-antibody TMT labeling^3,20^. The peptide flow-throughs of the UbiFast enrichment step containing unlabeled, non-K-ε-GG peptides are further processed for deep-scale and highly-multiplexed measurement of the proteome, phosphoproteome and acetylome using previously described methods.

Here we systematically evaluate each step of the serial MONTE workflow and apply the optimized method in a proof-of-concept study of primary patient lung adenocarcinoma (LUAD) tumors. The results demonstrate that the depth of coverage and quantitative precision at each of the ‘omes is not compromised by adding HLA peptidome and ubiquityl-peptide enrichments in serial with proteome, phosphoproteome and acetylome analysis. HLA immunopeptidomics of these pilot samples identified putative immunotherapeutic targets such as cancer/testis antigens and neoantigens. By enabling multiple, key ‘omes to be obtained on limiting amounts of exactly the same human tissue samples, MONTE overcomes prior limitations that have prevented concordant readout of the immunopeptidome, proteome and PTM-omes (ubiquitylome, phosphoproteome, acetylome) thereby enabling new insights into cancer and other disease biology.

## Results

### Native serial enrichment of the HLA immunopeptidome, proteome, and post-translational modifications by MONTE

To address the challenge of deeply characterizing clinically relevant samples with limited cellular input, we serialized HLA-I and HLA-II immunopeptidomics with ubiquitylome, proteome, phosphoproteome, and acetylome profiling workflows. The Multi-Omic Native Tissue Enrichment (MONTE) is represented in **Figure 1**. Three major changes were made to previously reported serial multi-omic enrichment protocols^3,4,13,14,16,17^ that were evaluated to ensure that each proteomic data type was not significantly biased. These changes included i) putting UbiFast K-ε-GG peptide enrichment before serial, multiplexed proteome, phosphoproteome, and acetylome analysis, ii) adding a protein level, serial HLA-II and HLA-I immunopeptidome enrichment prior to the downstream multi-omics analyses listed above, and iii) switching from an 8M urea cell lysis to an SDS cell lysis and digestion on an S-Trap to facilitate removal of detergents present in the native lysis buffer used for HLA IP. We also incorporated a modified version of a semi-automated, 96-well plate based HLA immunopeptidomics workflow ^25^ along with improved automated phosphopeptide enrichment^26^ and automated UbiFast K-ε-GG peptide enrichment^20^ workflows that enable higher throughput and reproducibility. Evaluation and optimization of each step of the MONTE workflow is detailed, below.

**Figure 1:**
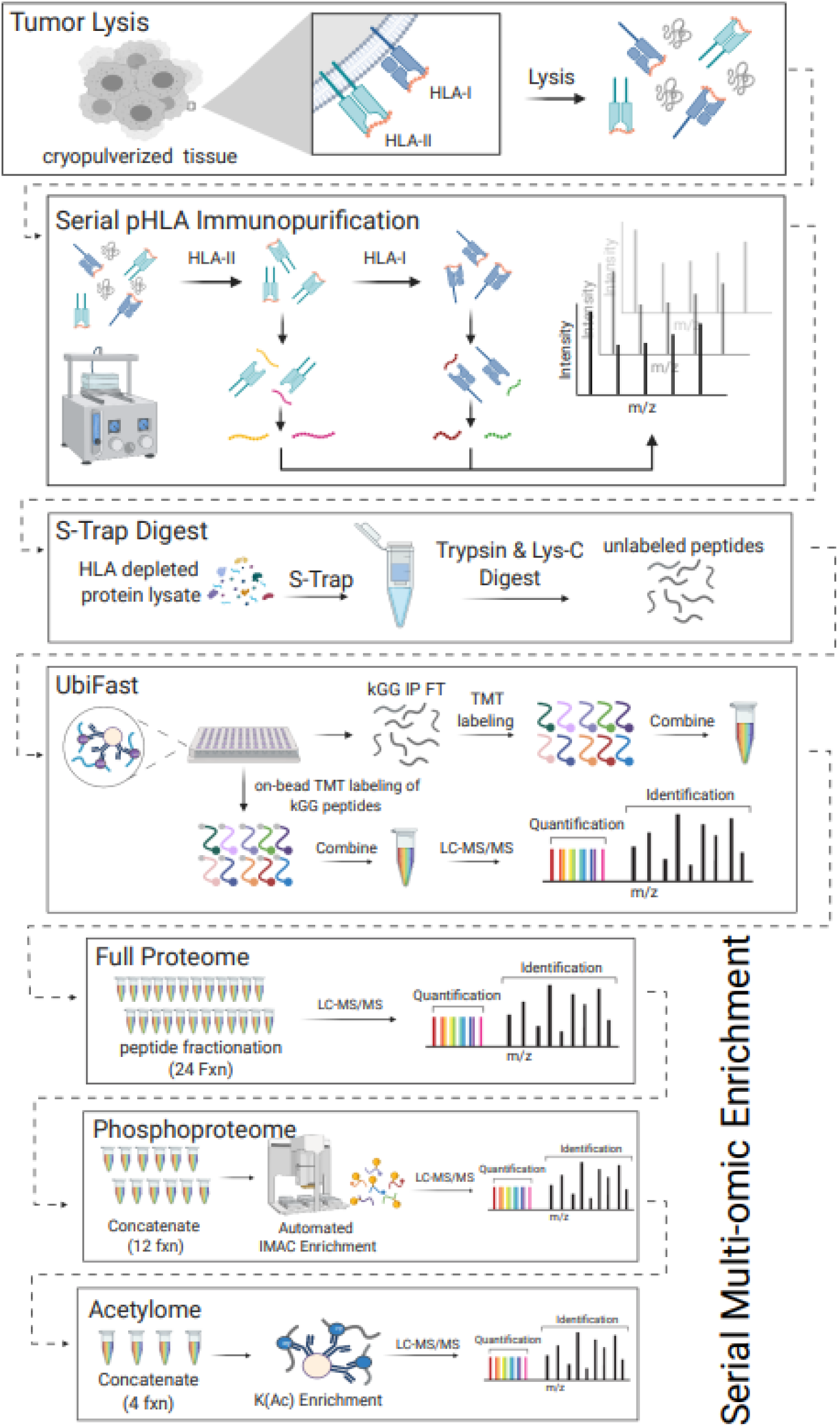
Schematic Overview of MONTE. The Multi-Omic Native Tissue Enrichments (MONTE) workflow for serial HLA immunopeptidome (label free), ubiquitylome (label free enrichment followed by TMT labeling post elution), proteome (TMT), phosphoproteome (TMT), and lysine acetylome (TMT). Dashed lines indicate use of the flow-through from each step.

### UbiFast followed by serial multi-omic sample processing shows expected coverage, reproducibility and retention of biological information

We first sought to integrate the UbiFast workflow with our well established TMT multiplexed proteome, phosphoproteome and acetylome workflow^4,16,17^. For this, we created the new workflow shown in **Figure 2A** that starts with the UbiFast method for enrichment and on-antibody TMT labeling of K-ε-GG peptide^3,20^. After UbiFast processing, flow-throughs from the antibody enrichment step that contain unlabeled, non-K-ε-GG peptides are subsequently TMT labeled and used as input to generate proteome, phosphoproteome and other PTM profiling datasets.

**Figure 2:**
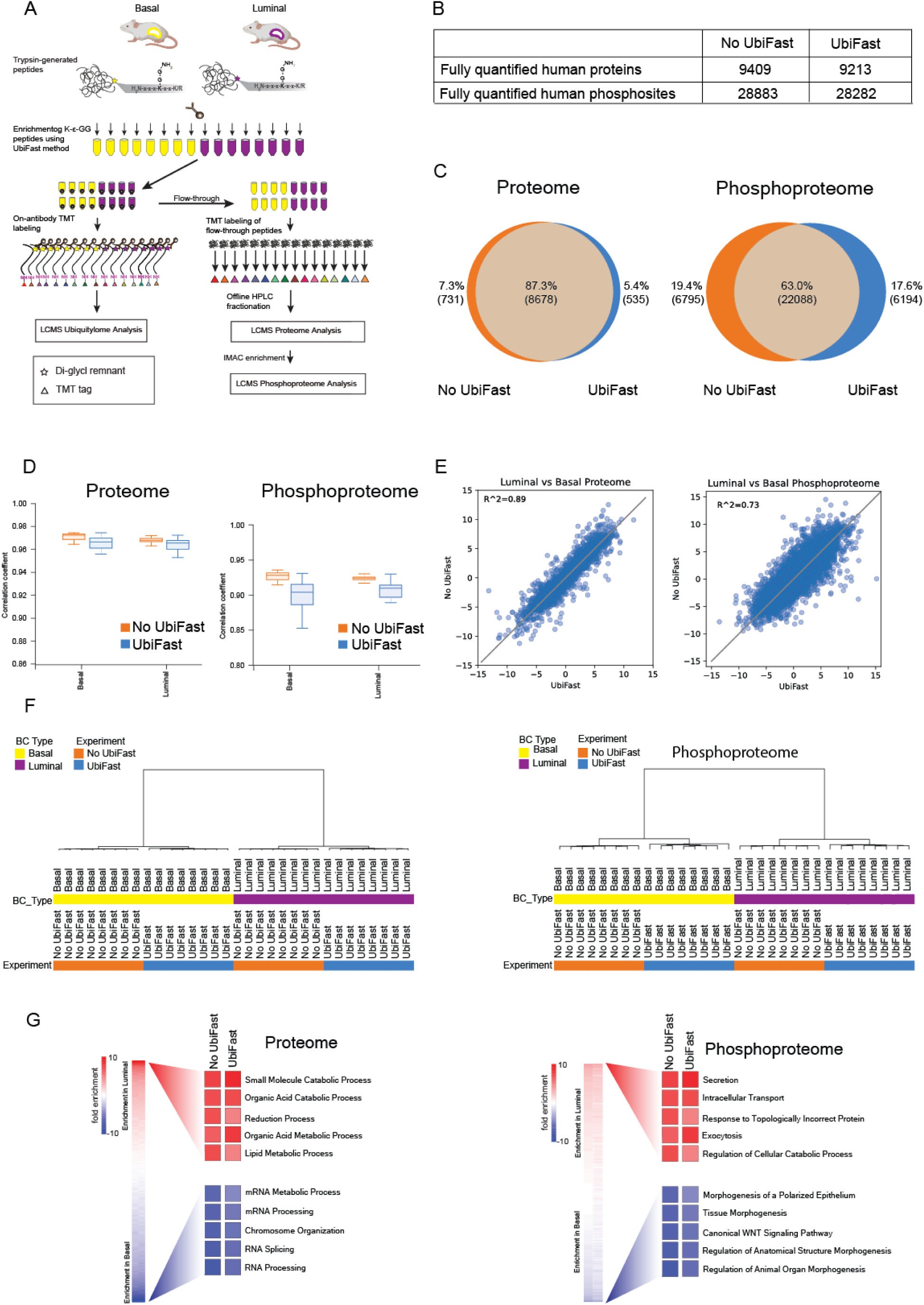
Comparison of the proteome and phosphoproteome of PDX samples with and without UbiFast pre-enrichment. **A**) Experimental design used to analyze K-ε-GG peptides, proteins and phosphopeptides from the same PDX samples.The flow-through of the UbiFast enrichment was used for proteome and phosphoproteome using 250 µg peptide input per TMT state. **B**) Summary table reporting the numbers of proteins and phosphorylation sites detected from non-UbiFast enriched (No UbiFast) and UbiFast enriched (UbiFast) CompRef PDX samples. **C**) Venn diagrams showing the overlap of proteins and phosphorylation sites for No UbiFast and UbiFast samples. **D**) Pearson correlation between Luminal and Basal PDX replicates (n = 8 luminal, n = 8 basal process replicates) for proteome and phosphoproteome data. Boxplots depict upper and lower quartiles, with the medians shown as a solid line. Whiskers show 1.5 interquartile range. **E**) Scatter plot showing log2 fold-change between Basal and Luminal PDXs, for proteins and phosphorylation sites. **F**) Hierarchical clustering for Proteome and Phosphoproteome separated by breast cancer subtype and +/- UbiFast. **G**) Heatmaps of all gene sets present in both experiments with the top and bottom 5 most enriched gene sets highlighted for both the proteome and phosphoproteome.

The addition of UbiFast was evaluated using tumors isolated from breast cancer patient-derived xenograft (PDX) models, representing Basal (WHIM2) and Luminal (WHIM16) subtypes of breast cancer^3,17,27^. We previously showed that we obtain >14,000 distinct Ub-peptides from these samples starting with 0.5 mg peptide input per channel^3,20^. Unlabeled peptide flow-throughs from KGG-Ab captures corresponding to 0.25 mg input peptide per state were subsequently labeled with TMT and combined for serial proteome and phosphoproteome analysis. Analysis by LC-MS/MS showed expected coverage of both the proteome and phosphoproteome with 9,165 human proteins and 28,406 human phosphorylation sites identified and quantified from the UbiFast flow-through samples(**Figure 2B**). The overlap of proteins and phosphorylation sites between experiments with and without serial UbiFast processing was high (87.3% proteome and 63.0% phosphoproteome)(**Figure 2C**). Pearson correlations of TMT ratios between intraplex replicates for UbiFast flow-throughs were high with median correlations of 0.97 for both Basal and Luminal subtypes, indicating that UbiFast pre-processing does not negatively affect reproducibility(**Figure 2D**). Scatter plots of Basal/Luminal protein and phosphosite TMT ratios measured in UbiFast flow-through samples and non-UbiFast samples correlated well (R^2 = 0.89 proteome and R^2 = 0.73 phosphoproteomes)(**Figure 2E**). Unsupervised hierarchical clustering of proteome and phosphoproteome samples show a dramatic separation of samples by breast cancer subtype as expected with much smaller separation by experiment(**Figure 2F**). Finally, pathway enrichment of regulated proteins and phosphorylation sites correlated well with the top regulated gene sets in data acquired without pre-UbiFast processing(**Figure 2G**). Taken together, these results support the feasibility of incorporating multiplexed ubiquitylation profiling using UbiFast up-front and serially with multiplexed proteome and PTM profiling workflows.

### Addition of serial HLA-II and HLA-I enrichment into the MONTE workflow achieves similar data depth to analyses done in parallel

We evaluated the impact of adding serial HLA-II and HLA-I enrichment prior to serial multi-omic enrichment workflow for ubiquitylome, proteome, phosphoproteome and acetylome. For these studies we used ten cryopreserved primary lung adenocarcinoma (LUAD) tumors from the Clinical Proteomic Tumor Analysis Consortium (CPTAC) cohort^12^. Human tumor samples are more relevant than tumor samples derived from immunocompromised mice, and in testing, we found that the yield of HLA-I and HLA-II immunopeptidomes from the PDX breast cancer tumor models was too low to derive meaningful conclusions. LUAD tumors were selected because lung tissue is known to have HLA-I and HLA-II expression^28,29^ and one LUAD primary tumor has been profiled successfully using serial HLA-I and HLA-II immunopeptidomics^22^. This set of LUAD samples was chosen to represent important biological differences of high relevance to lung adenocarcinoma, as five samples were driven by mutant KRAS and five by EGFR mutations (four of which were in never-smokers). None of these previously characterized tumors were from the immune hot cluster^12^, and each driver mutation subset included samples from both men and women and both Asian and Western/Caucasian ethnicity were represented. The human LUAD tumors (50-86 mg cryopulverized tissue) were lysed in SDS and processed with and without initial serial HLA enrichment(**Figure 3A)**. S-Trap-based protein digestion^24^ was used instead of 8M urea digestion post HLA enrichment because we have previously shown that serial HLA immunopeptidome and downstream whole proteome analysis required the removal of detergents present in the native lysis buffer used for HLA enrichment^30^.

**Figure 3:**
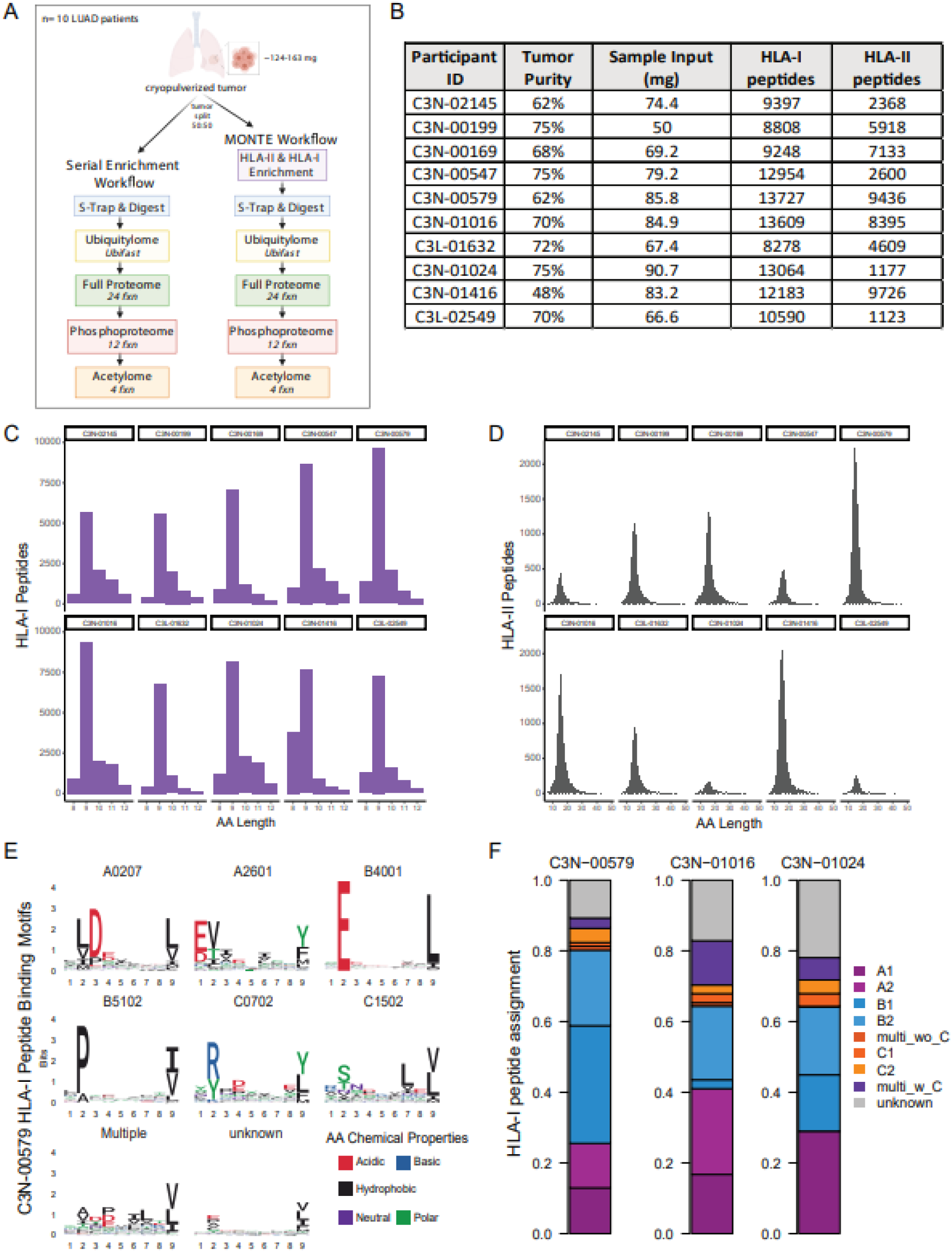
LUAD tumor HLA-I and HLA-II immunopeptidome profiling using MONTE. **A**) Schematic overview of the head-to-head serial multi-omic enrichment vs. MONTE workflows used to characterize ten cryopulverized primary LUAD tumors. **B**) Table summarizing the tumor input and resulting HLA-I and HLA-II peptides mapping to human source proteins detected from ten LUAD patients. **C**) Length distributions of HLA-I peptides from the LUAD cohort. **D**) Length distributions of HLA-II peptides from the LUAD cohort. **E**) Example HLA-I peptide motifs and the allele expressed by LUAD Patient C3N-00579. **F**) Example HLA-I peptide:allele assignments obtained using the presentation predictions from HLAthena for three LUAD patients^7,8^.

Label free, antibody-based, serial HLA enrichment identified a median of 11,387 HLA-I (8,278-13,727) and 5,263 HLA-II (1,123-9,726) bound peptides from each of these ten LUAD tumors(**Figure 3B**). Our depth of >10,000 HLA-I peptides from as little as 50 mg cryopulverized tumor corresponding to ∼2 mg protein lysate was encouraging, and clearly indicated that the method would likely be usable with even smaller amounts of input tumor material. We confirmed that the observed HLA-I and HLA-II peptides had the expected length distributions(**Figure 3C, D**) and HLA-I binding characteristics(**Figure 3E, F**) using a motif analysis and the HLA-I presentation predictor HLAthena^7,8^. Patient C3N-01416 had a larger representation of 8mers in the HLA-I immunopeptidome, which was expected because of the known preference for 8mers presented by HLA-B*18 alleles. We also confirmed that HLA-II immunopeptidomes contained motifs consistent with patient HLA-II alleles called from RNAseq data by arcasHLA^31^(**Figure S1**).

The protein flow-throughs from HLA immunopeptidome enrichments were next digested with Lys-C and trypsin using S-Traps in parallel with half of each LUAD tumor that was not HLA enriched. A summary of the resulting depths of these head-to-head proteomes, ubiquitylomes, phosphoproteomes, and acetylomes are shown in **Figure 4A**. The proteome and ubiquitylome results demonstrate that similar numbers of canonical human proteins (11,028 vs.10,729) and K-ε-GG peptides (9,516 vs. 9,419) were identified and fully quantified between the non-HLA enriched (“No HLA”) and HLA enriched (“HLA FT”) samples, respectively. A 16% decrease in the total number of phospho-sites (−8% phosphorylated proteins) was observed when using the HLA enriched samples (No HLA: 26,627 phosphosites, 6,745 phosphoproteins; HLA FT: 22,339 phosphosites, 6,235 phosphoproteins), suggesting that the phosphatase inhibitors added to our lysis buffer may be losing their activity during the protein-level, HLA immunopeptidome enrichment. The number of lysine residues observed to be acetylated on internal lysine residues (i.e, not at the N- or C-termini or the peptide) increased by 45% in the HLA enriched samples (No HLA: 3,702 vs. HLA FT: 5,280 internal K-acetylsites). The relative yield of acetylated peptides (i.e., relative yield is the percentage of K-Ac peptides relative to the total peptides identified in the sample) in the HLA processed samples was significantly higher (75% vs. 55%). Given that the protein lysates were incubated at 4°C for 6 hr during HLA enrichment, we sought to rule out possible non-enzymatic acetylation^32^. Acetylome analysis of A375 melanoma cells with and without the 6h HLA IP incubation conditions yielded a similar number of acetylated peptides when compared to no HLA incubation conditions, suggesting the addition of the HLA IP did not cause non-enzymatic acetylation. Another possible explanation for increased acetylation sites could be due to pre-clearing of non-specifically binding components in the complex tissue lysates by HLA- and K-ε-GG antibodies, resulting in increased relative yield.

**Figure 4:**
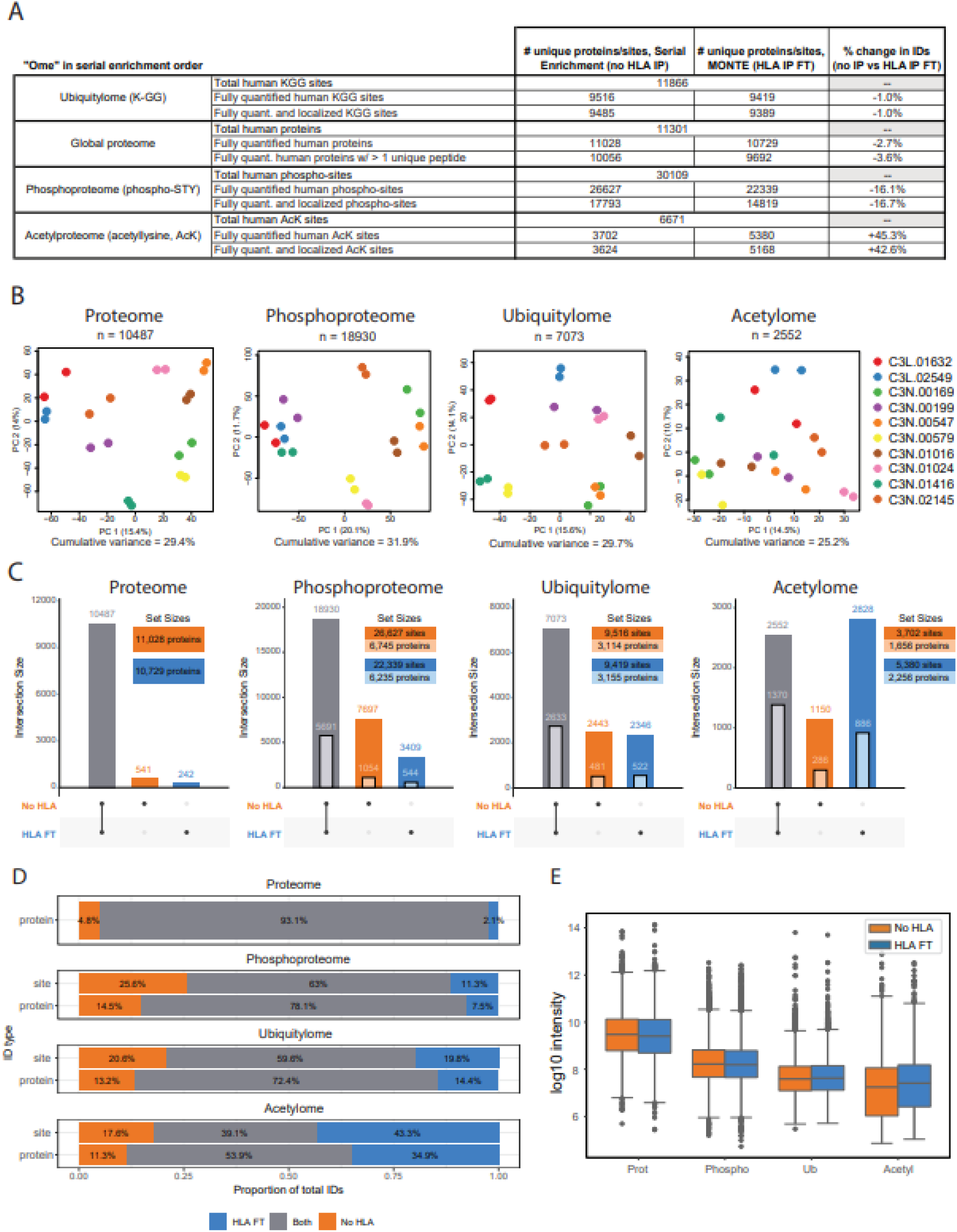
Evaluation of data depth between serial multi-omic enrichment with and without HLA enrichment. **A**) Summary table reporting the numbers of proteins and PTM containing peptides detected from non HLA enriched (No HLA) and HLA enriched (HLA FT) LUAD samples. **B**) Principal component analysis of LUAD tumors (n=10) for all human proteins and PTM sites quantified in all TMT channels in both No HLA and HLA FT conditions. **C**) The total number of quantified human proteins and PTM site identifications illustrated as ‘UpSet’ plots^76^. Vertical bars depict the number of uniquely or jointly detected features, as indicated by the layout matrix below. Darker colored bars represent unique proteins in the proteome and PTM sites in each PTM-ome. Lighter colored nested bars represent the number of unique modified proteins in each PTM-ome. **D**) Stacked bar chart showing proportional overlap for No HLA and HLA FT conditions by unique proteins in the proteome and by PTM sites or modified proteins for each PTM-ome. **E**) Log10 total intensity distributions of all human proteins and PTM sites from No HLA and HLA FT samples. Boxplot depicts upper and lower quartiles, with the median shown as a solid line. Whiskers show 1.5 x interquartile range.

### Biological signals observed in a serial multi-ome enrichment workflow without HLA enrichment are recapitulated in the MONTE workflow

To assess potential differences between HLA enriched and non HLA enriched samples, we analyzed the ten LUAD tumor proteomes, ubiquitylomes, phosphoproteomes, and acetylomes using a principal component analysis(PCA)(**Figure 4B**). PCA analysis shows that samples cluster by LUAD tumor, not by the processing method used, demonstrating that biological differences among the samples are stronger than technical variation between these serial workflows. The acetylomes appeared to cluster the least, likely due to the increase in acetylated peptide yield observed when using the flow-through from the HLA enrichment step. The total number of proteins identified and quantified from HLA enriched and non-HLA enriched samples were shown to have an 93% overlap(**Figure 4C, D**). Slightly fewer proteins (3%) were identified from the HLA enrichment flow-throughs due to the depletion of HLA complexes and associated binding partners, including a minor depletion in the proteins involved in the microtubule organizing center (MTOC). This may relate to the direct binding of MTOC proteins with HLA peptide complexes embedded in membranes, as the HLA-II presentation pathway requires endocytic compartments that contain motor proteins like dynein and kinesin^33,34^.

The overlap between HLA enriched and non enriched protein lysates was 60% for ubiquitylation sites, 72% for ubiquitylated proteins, 63% for phosphorylation sites and 78% for phosphorylated proteins, which is an expected result using multiplexed, data-dependent LC-MS/MS methods for highly similar processing workflows^17^(**Figure 4C, D**). The lowest overlap across experiments was observed for acetylome data because 45% more acetylated peptides were observed in the HLA enriched samples. Overall, the HLA enriched samples capture the same depth of coverage observed in non-HLA enriched samples and adding this enrichment step upfront in a serial workflow does not introduce detectable bias in downstream proteome, ubiquitylome, phosphoproteome, and acetylome.

### HLA peptides from mutated, noncanonical, and cancer-testis antigen source proteins are identified in MONTE immunopeptidomes

MONTE immunopeptidomes were analyzed using a personalized database containing canonical human proteins, noncanonical proteins from novel or unannotated open reading frames (nuORFs), mutations shared across TCGA tumor types, and patient specific mutations(**Figure 5A**). Initially, we looked in the LUAD immunopeptidomes for peptides derived from cancer-testis antigen (CTA) source proteins reported in the CTdatabase and observed peptides from 45 unique source proteins^35,36^. Across the set of LUAD tumors, peptides derived from seven CTA source proteins specific to lung cancers^36^ were detected, including two from the MAGE family(**Figure 5B**). Most peptides from CTA source proteins were presented by HLA-I with the exception of TEXT101 and ACTL8 presented by HLA-II. Surprisingly, two unique peptides derived from the bromodomain testis-specific protein (BRDT) were presented by 6/10 our LUAD tumor set, making it a candidate for future immunogenicity investigations.

**Figure 5:**
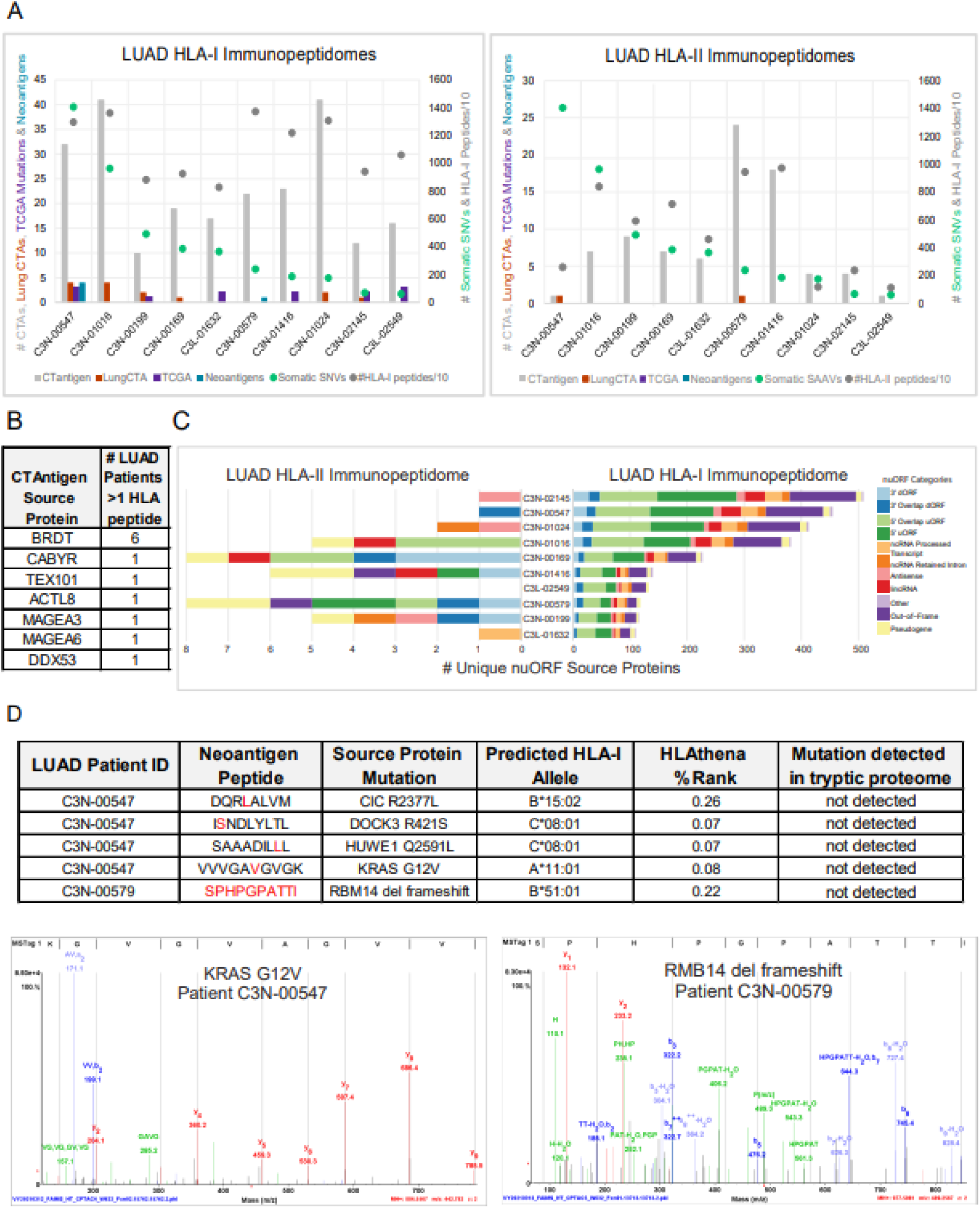
Analysis of HLA peptides presented by LUAD tumors derived from CT antigen, nuORF, and mutation containing source proteins. **A)** Summary of unique HLA-I and HLA-II peptides (dark gray), total somatic single nucleotide variants (SNVs; green), peptides mapping to source proteins from the CTdatabase (light gray), peptides mapping to lung cancer specific^36^ CTA source proteins (orange), peptides containing TCGA shared mutations (purple), and neoantigens containing patient specific somatic mutations (blue) from ten LUAD tumors. The primary y-axis shows counts of CTAs, Lung CTAs, peptides containing TCGA shared mutations, and neoantigens. The secondary y-axis shows the total number of somatic SNVs and the immunopeptidome depths per LUAD patient. **B**) A summary table reporting the number of LUAD patients presenting at least one HLA-I or HLA-II peptide from CTA source proteins reported in^36^. **C**) A mirrored stacked bar plot showing the number of unique nuORF source proteins that are presented by HLA-I (right) and HLA-II (left) colored by the nuORF category^37^ across ten LUAD patients. **D**) Summary table of HLA-I neoantigens detected in primary LUAD patient tumors (top) accompanied by annotated MS/MS spectra for KRAS G12V (bottom, left) and the RBM14 frameshift (bottom, right) neoantigens. See Supplemental Information for annotated MS/MS spectra of the others.

Next, we sought to detect peptides in our LUAD HLA immunopeptidomes derived from nuORFs whose translation has been supported by ribosome profiling using a recently published nuORF database^37^(**Figure 5C**). High-confidence HLA-I and HLA-II peptide identifications derived from nuORFs were found across 9/10 patients (see **Methods**). HLA-I immunopeptidomes contained far more unique nuORF source proteins than HLA-II, and the overall ranking of patients by number of unique nuORF source proteins did not correlate between HLA-I and HLA-II immunopeptidomes. The average representation of nuORF source protein categories per sample also differed between HLA-I and HLA-II, as a higher proportion of HLA-II nuORFs mapped to pseudogenes (19%) and few mapped to out-of-frame ORFs (5%), while the reverse was true for HLA-I where the total percentage of pseudogenes and out-of-frame ORFs were 3% and 21%, respectively. These observations align with recent studies suggesting that the HLA-I pathway is more likely to sample less stable, shorter proteins, while the HLA-II pathway is more likely to sample stable source proteins^22,38^. The contrasting nuORF representations also highlight the differences in noncanonical source protein presentation between HLA-I and HLA-II pathways that are not yet fully understood but could be improved upon from data obtained on larger patient cohorts.

We then assessed if the LUAD immunopeptidome depth enables the detection of HLA peptides containing patient specific mutations. Historically, detection of neoantigens by LC-MS/MS has required enrichment from either billions of cells or gram-levels of tissue as neoantigens are presented at levels as low as 0.01%^2,39,40^. We hypothesized that recent improvements in MS instrumentation, off-line fractionation, and separation in the gas phase^23^ could enable detection of neoantigens from the much smaller amounts of input sample. We first mined our data for HLA peptides containing mutations shared across multiple tumor types reported in TCGA^41^. Six of the ten patients had at least one HLA-I peptide mapping to a shared TCGA mutation, while no HLA-II peptides containing TCGA mutations were identified. The most frequently presented TCGA mutation was a point mutation (H33Q) in IDUA, an enzyme found in lysosomes, that was presented by three of the ten patients on either HLA-B*55:02 or -B*37:01. We also observed HLA-I peptides containing TCGA mutations presented by at least two of the patients from SEMA3A (point mutation) and FCGBP (point mutation). By comparing published whole exome sequencing (WES) data from both blood (normal) and tumor from these LUAD patients^12^, we determined that the IDUA mutation (10/10 patients) was germline and not a neoantigen. Interestingly, several of the other TCGA shared mutations found in HLA-I peptides, such as those from ATP6AP2, SEMA3A, and FCGBP were not called in any patient by the WES data and will require further validation before they can be confirmed as putative neoantigens.

To find HLA presented neoantigens, we analyzed the immunopeptidomes for peptides containing somatic mutations^42–44^. Two of the ten patients (20%) had at least one detected neoantigen in their HLA-I immunopeptidomes, of which four contained point mutations and one a frameshift deletion mutation(**Figure 5D, Figure S2**). Most neoantigens were derived from mutations not shared across patient populations with the notable exception of the KRAS G12V neoantigen detected in patient C3N-00547. The KRAS G12V 10mer is a shared neoantigen that has been previously confirmed to be presented on HLA-A11^45,46^, and to our knowledge, this is the first report of LC-MS/MS detection from a primary patient LUAD tumor. We also detected two neoantigens bound to the less abundantly expressed HLA-C alleles, perhaps aided by the very similar binding specificity of the patient’s two alleles C*08:01, C*12:03. In general, we observed that patients with high mutation burden and immunopeptidome depth(>10,000 peptides) were most likely to have LC-MS/MS detectable neoantigens. These results suggest that detection of neoantigens by immunopeptidomics should, at present, be focused on tumor types with relatively high mutational burden, high HLA expression levels, and only the most highly optimized LC-MS/MS methods should be used.

### Integrating immunopeptidomics with whole proteome and PTM-omes reveals insights into source protein processing and presentation

An advantage of the MONTE workflow is that the resulting multi-omic data is derived from each single sample, enabling robust data integration. Thus, we evaluated whether integration of MONTE data would capture known as well as new biological insights. We first looked at how well HLA-I and HLA-II source proteins overlapped with both the proteome and ubiquitylome data (**Figure 6A**). 78% (9274/11831) of proteins identified in the proteome were also identified as HLA-I source proteins, while 30% (4011/13285) of HLA-I source proteins were not observed in the proteome. Alternatively, a 33% (3871/11831) overlap between the proteome and HLA-II source proteins was observed with 21% (1059/4930) of HLA-II source proteins not detected in the proteome, which is likely due to differences in source protein sampling bias between the HLA-I and HLA-II pathways. A higher proportion of ubiquitylated proteins, 89% (3499/3954), were detected as HLA-I source proteins, compared to 49% (1939/3954) as HLA-II source proteins. This was expected because ubiquitylated proteins are a key source of proteasomal processed peptides that are HLA-I peptide precursors. Conversely, we noted only 26% (3499/13285) of HLA-I source proteins were identified as ubiquitylated, suggesting that deeper ubiquitylome datasets are required to fully overlap with HLA-I immunopeptidomes. Because HLA-I source protein expression levels and their ability to be processed by the proteasomal pathway are important factors for presentability, both proteome and ubiquitylome datasets are useful for incorporation into HLA-I prediction algorithms.

**Figure 6:**
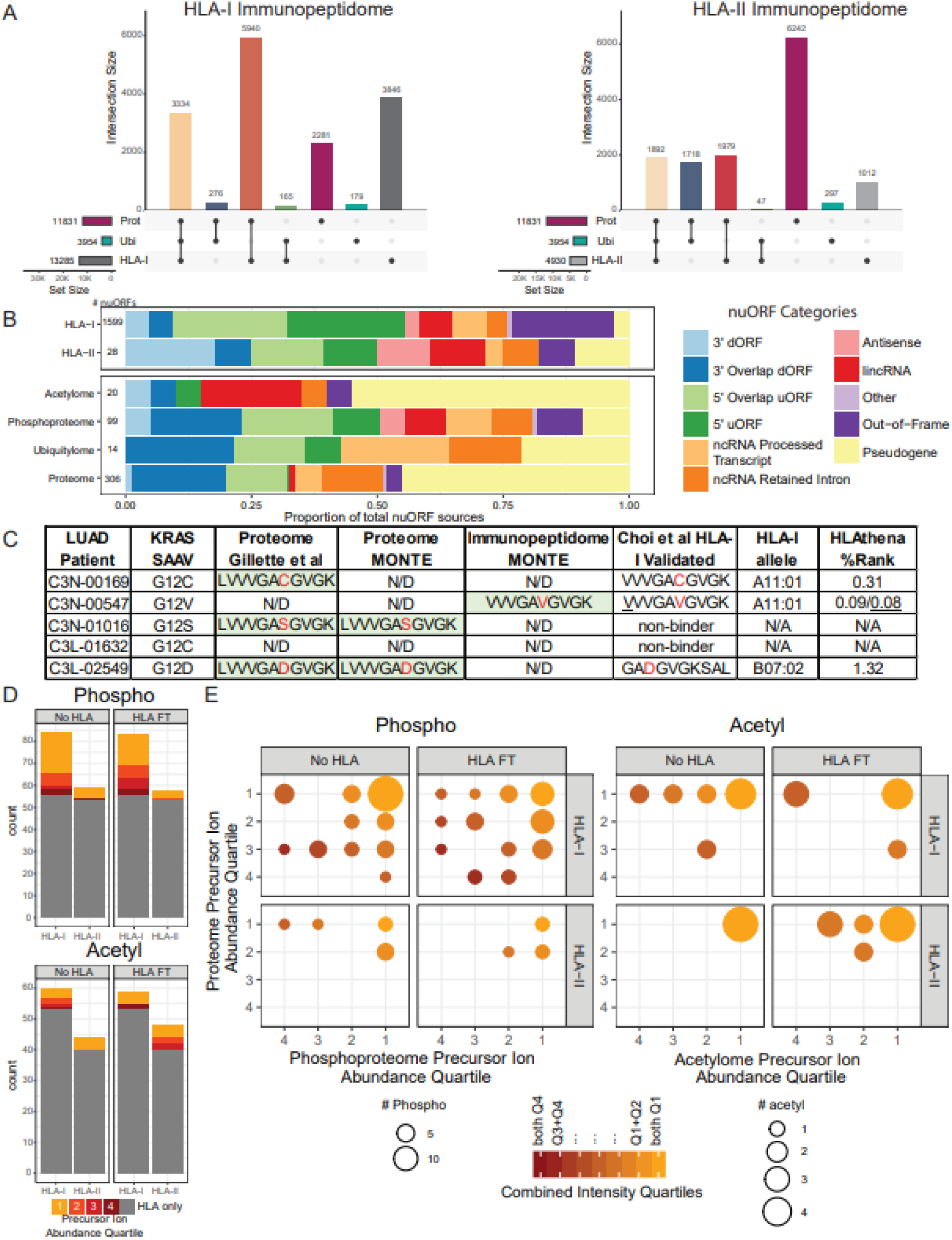
Integration of LUAD MONTE immunopeptidome data with whole proteome and PTM-omes. **A**) The total number of proteins detected in proteome, ubiquitylome (ubiquityl-proteins), and immunopeptidome (source proteins) illustrated as ‘UpSet’ plots^76^. Vertical bars depict the number of uniquely or jointly detected features, as indicated by the layout matrix below. Color of vertical bars corresponds to each combination of sets. **B**) Stacked bar chart showing proportion of detected nuORF source proteins by nuORF type across the immunopeptidomes, proteomes, and PTM-omes. **C**) Table of patient-specific KRAS mutations and their detection in different datasets, including: tryptic proteome of large cohort study of LUAD^12^; MONTE tryptic proteome; MONTE HLA-I immunopeptidome. KRAS neoantigens previously identified by LC-MS/MS^45^ are shown as well as predictions for presentation for the best HLA-I allele using HLAthena^8^. The wildtype KRAS G12 peptide was not detected in the MONTE proteomes of this small subset of the LUAD patient samples with or without initial HLA IP), but it was detected in 23/25 LUAD TMT10 plexes covering 102/111 patients in the original study^12^. **D)** Stacked bar charts showing overlap of phosphorylated and acetylated immunopeptides with sites detected and unambiguously localized in the phospho- and acetyl-proteome. Color indicates precursor ion intensity quartile of detected modification in the PTM-ome (1 = highest, 4 = lowest), with gray indicating that the site was uniquely detected in the immunopeptidome. **E**) Marble plots show relative abundance of PTM-sites identified in the immunopeptidome as detected in the phospho- or acetyl-proteome, as well as the abundance of corresponding phospho- or acetyl-proteins as detected in the proteome. Size of points encodes number of sites and color encodes combination of precursor intensity quartiles of PTM-ome and proteome.

We also investigated the representation of nuORFs in the MONTE proteomes and PTM-omes compared to those detected in the immunopeptidomes(**Figure 6B**) and observed that the representation of nuORF categories in HLA-I immunopeptidomes and phosphoproteomes were the most diverse compared to all other ‘omes. In general, a higher proportion of pseudogenes were detected in the proteome and PTM-omes when compared to HLA-I and HLA-II immunopeptidomes. We also noted that the representation of nuORF categories varied across the different ‘omes, with the acetylome and ubiquitylome having the highest proportion of lincRNAs and noncanonical RNA processed transcripts, respectively, while the HLA-I immunopeptidome contained the most out-of-frame ORFs. The HLA-I immunopeptidome yielded detection of greater than 5 times more nuORFs than any other ‘ome. Hence, while many nuORFs are translated and may be capable of becoming antigens, some are post-translationally modified and therefore may be involved in regulating cellular pathways.

We next asked if mutations resulting in LC-MS/MS detectable neoantigens were present in MONTE proteomes. None of the mutations contained within detected HLA-I neoantigens (**Figure 5D**) were detected within tryptic peptides from the proteome. Given that 5/10 of the LUAD patient samples analyzed carry a KRAS G12X mutation that is contained within the tryptic peptide LVVVGAXGVGK, we examined the overlap in KRAS mutation detection between the immunopeptidome and proteome(**Figure 6C**). We noted that three patients with KRAS G12V/D/C mutations (C3N-00169, C3N-00547 and C3L-02549) expressed HLA-I alleles that have been validated to present KRAS neoantigens^45,46^. Only the KRAS G12V 10mer was detected in HLA-A11 homozygous patient C3N-00547, which is more likely than the G12C and G12D variants to be presented (HLAthena %rank 0.08 vs. 0.31 and 1.32, respectively). Although our MONTE immunopeptidomes were not able to capture all validated KRAS neoantigens found using an artificial overexpression system^45^, this limitation will diminish as even more sensitive MS instrumentation and data generation approaches are introduced. Surprisingly, tryptic peptides containing the KRAS G12V mutation were not detected by LC-MS/MS in the MONTE proteome data or in our earlier 111 patient LUAD study from which the ten patient samples analyzed here were obtained^12^. Overall, 32/111 patients in the study had KRAS G12 mutations, of which only G12C, G12D, and G12S were detected as tryptic peptides (SAAV/SNV: 4/16, 2/7, and 2/2 patients respectively), while G12A and G12V were not detected (SAAV/SNV: 0/2 and 0/5 patients respectively). The lack of KRAS G12V tryptic peptide detection in the proteome demonstrates that low source protein expression was overcome by strong HLA-I binding and stability resulting in neoantigen detection.

The immunopeptidomes were also searched for PTM modified peptides. We observed HLA-I and HLA-II phosphopeptides at 0.11% and 0.3%, respectively, and acetylpeptides at 0.08% and 0.10%, respectively, of total unique peptides. Prior studies have shown that position four in HLA-I peptides is the residue most often phosphorylated^47–49^. Consistent with these studies, we find a majority of HLA-I phosphorylation sites (54%) on the fourth amino acid. However, acetylation appeared only 10% in the fourth position and more often in the first position (31%). As expected, HLA-I phosphopeptide detection was more likely to occur in patients with HLA-I alleles that contain proline in their binding motifs that correspond to the kinase substrate motifs of MAPK and CDK^47,50^. We next evaluated if the abundance of phospho- or acetyl-sites(**Figure 6D**) and corresponding phospho- or acetyl-proteins detected in the proteome(**Figure 6E**) impacts HLA-I and HLA-II peptide presentation. While many of the PTM containing source proteins were also observed in their corresponding PTM-ome (HLA-I: phospho 87%, acetyl 50%; HLA-II: phospho 42%, acetyl 43%), we observed that few of the specific PTM sites presented by HLA were detected in phosphoproteomes and acetylomes (HLA-I: phospho 42%, acetyl 13%; HLA-II: phospho 10%, acetyl 17%). Of the PTM sites that were observed in both the immunopeptidome and corresponding PTM-ome, we found that PTM containing source proteins in the top abundance quartiles are most likely to result in HLA presentation. Thus, source protein abundance is a factor that governs PTM HLA peptide presentation and may be useful for future PTM HLA peptide prediction efforts. However, confirmation of these preliminary observations will require focused analysis of larger datasets.

## Discussion

Patient tissue samples with limited input amounts create obstacles for deep proteomic characterization. MONTE directly addressed this limitation, enabling serial HLA-I and HLA-II immunopeptidome profiling followed by ubiquitylome, proteome, phosphoproteome, and acetylome analysis of each single sample. Two technical hurdles had to be addressed to fully serialize MONTE. First, immunopeptidomics was performed prior to the serial proteome and PTM-ome enrichment workflow, which required native lysis conditions with intact proteins captured and digested on S-Traps. Second, the enrichment of K-ε-GG peptides by UbiFast was completed before TMT labeling of peptides and other PTM enrichments because the anti-K-ε-GG antibody will not capture TMT-GG-ε-K. After implementing these changes to the serial workflow, we observed high correlation between the proteomes, ubiquitylomes, phosphoproteomes and acetylomes in both our breast cancer xenograft and LUAD datasets, and we were able to recapitulate expected biological signals, demonstrating that putting HLA and UbiFast enrichment upfront of the serial enrichment workflow does not negatively impact downstream proteome and PTM-ome data.

In proteogenomic studies to date, HLA immunopeptidomics has not been routinely employed and ubiquitylomics is only now beginning to be used to profile human tumor samples^15^. Layering of HLA-I and HLA-II immunopeptidomes on these other data types provides a window into the antigen landscape and improves our understanding of the rules that govern antigen processing and presentation. For example, patient C3N-00169 had a truncation mutation, E269*, in the proteasomal subunit PSMB7. We noted this patient was homozygous for HLA-A11 alleles that have a lysine residue in the C-terminal anchor position. This observation suggests that tryptic proteasomal subunits like PSMB7 may be under selection pressure in patients with HLA-I alleles that favor tryptic-like peptides. Furthermore, immunopeptidome and proteome datasets from the same sample could enable more accurate neoantigen and noncanonical HLA peptide prediction methods, as having both HLA presentation and protein expression data can be used to improve epitope prediction algorithms^7,8,51–54^.

As noted with the KRAS G12V neoantigen and nuORF derived HLA peptides, tryptic proteome detection alone, if used for neoantigen and non-canonical peptide prediction, would likely under-represent the full neoantigen and noncanonical peptide repertoires. As such, MONTE immunopeptidome and proteome datasets from larger cohorts are required to fully understand how best to integrate tryptic proteome level mutation detection into epitope prediction workflows. Similarly, PTM-ome data combined with immunopeptidomics can uncover dependencies, such as PTM abundance, that can be used to improve prediction of difficult to detect antigens containing PTMs. Integrated MONTE datasets may also provide information regarding tumor immune cell infiltration status and dysregulation of signaling, degradation, and epigenetic pathways that can inform therapeutic intervention.

High-throughput multi-omic data generation has proven to be a useful resource for understanding disease biology and the identification of therapeutic targets^11,14,55–62^. By combining serial multi-ome enrichments with HLA-I and HLA-II immunopeptidomics into a single workflow, we have provided a path forward to understanding connections between antigen presentation and protein expression, signaling, protein degradation, and epigenetic regulation from each single sample, which was not previously possible. Further improvements to the MONTE workflow that address its current limitations will likely include decreasing the sample input further by incorporating low-input proteomic sample processing advances^63–66^ and on-line, gas-phase fractionation technologies such as ion mobility^67,68^, and the incorporation of fully automated sample processing steps for all ‘omes. We also anticipate that additional ‘omes could be incorporated into MONTE such as phosphotyrosine peptide enrichment. In addition to cancer biology, the MONTE workflow is also applicable to the study of other disease states, such as autoimmune and infectious diseases to address questions related to disease pathology and treatment.

## Methods

### Serial Processing and Analysis of Proteome and Phosphoproteome from UbiFast flow-through samples of Comparative Reference Tissue(CompRef)

WHIM2 and WHIM16 patient-derived xenografts (PDX) were processed for proteomic analysis as described previously^17^. Briefly, frozen tissue from each model was lysed, digested and aliquoted into 0.5 mg aliquots. For this experiment approximately 20–50 mg of xenograft tissue was used to obtain 0.5 mg of peptide input. Enrichment of K-ε-GG peptides using 0.5 mg of peptide per sample was performed using the automated UbiFast method^3,20^ and non-K-ε-GG peptides were collected in the flow-through for proteome and phosphoproteome analysis.

For proteome analysis, peptides were acidified and desalted using a 50mg tC18 SepPak cartridge. Eluates were frozen and a vacuum centrifuge was used to dry peptides. Peptides were reconstituted in 30% ACN, peptide concentration was determined using a BCA assay and peptides were dried again. Peptides corresponding to 0.25 mg from each sample (eight replicates of WHIM2 and eight replicates of WHIM16), were labeled with 0.5 mg TMTPro reagent in 20% ACN, 50 mM HEPES for 1 hour. The TMT labeling reaction was quenched by adding 4 µL 5% hydroxylamine for 15 min at room temperature while shaking. Samples were combined into a 15 mL conical tube, frozen at -80°C and dried in a vacuum centrifuge. The combined sample was desalted using a 200 mg tC18 SepPak cartridge and the eluate was snap frozen then dried in a vacuum centrifuge. Offline bRP fractionation was performed as previously described ^17^. Briefly, peptides were separated over a 96 min gradient with a flow rate of 1 ml/min. The bRP solvent A was 5 mM ammonium formate, 2% ACN and solvent B was 5 mM ammonium formate, 90% ACN. 96 fractions were concatenated into 24 fractions for proteome analysis. For proteome analysis, 5% of each of the 24 fractions were transferred into HPLC vials, frozen and dried in a vacuum centrifuge.

The remaining 95% of each bRP fraction was concatenated into 13 fractions for phosphopeptide enrichment by IMAC. Concatenated fractions were reconstituted to a final concentration of 80% ACN / 0.1% TFA and phosphopeptides were enriched using the the Agilent “AssayMAP Phosphopeptide Enrichment v2.1” protocol on an Agilent Bravo system. Briefly, 200 µL of sample was loaded onto AssayMap Fe(III)-NTA cartridges (Agilent, G5496-60085) at 5 uL/min. The flow-through was collected and frozen for downstream acetyllysine enrichment. The cartridges were washed 3x with 80% ACN / 0.1% TFA and phosphopeptides were eluted from the cartridges with 20 µL fresh 1% ammonium hydroxide into a plate containing 2.5 µL neat FA. Phosphopeptides were transferred to HPLC vials, frozen and dried in a vacuum centrifuge. For LC/MS-MS analysis, peptides were reconstituted in 9 µL 3% ACN / 0.1% FA and 4 µL were injected from each of the 12 fractions.

### CompRef LC-MS/MS Analysis

All peptide samples were separated on an online nanoflow EASY-nLC 1200 UHPLC system (Thermo Fisher Scientific) and analyzed on an Orbitrap Exploris 480 mass spectrometer(Thermo Fisher Scientific). 1µg of each proteome and fifty percent of each phosphopeptide and K(GG) peptide sample was injected onto a capillary column (Picofrit with 10 µm tip opening / 75 µm diameter, New Objective, PF360-75-10-N-5) packed in-house with 25 cm C18 silica material (1.9 µm ReproSil-Pur C18-AQ medium, Dr. Maisch GmbH, r119.aq). The UHPLC setup was connected with a custom-fit microadapting tee (360 µm, IDEX Health & Science, UH-753), and capillary columns were heated to 50 °C in column heater sleeves (PhoenixST) to reduce back pressure during UHPLC separation. For proteome and phosphoproteome samples, injected peptides were separated at a flow rate of 200 nL/min with a linear 85 min gradient from 100% solvent A (3% acetonitrile, 0.1% formic acid) to 30% solvent B (90% acetonitrile, 0.1% formic acid), followed by a linear 10 min gradient from 30% solvent B to 90% solvent B. For ubiquitin samples, injected peptides were separated at a flow rate of 200 nL/min with a linear 120 min gradient from 100% solvent A (3% acetonitrile, 0.1% formic acid) to 35% solvent B (90% acetonitrile, 0.1% formic acid), followed by a linear 10 min gradient from 35% solvent B to 90% solvent B. Data-dependent acquisition was obtained using Xcalibur 4.4 software in positive ion mode at a spray voltage of 1.80 kV. MS1 Spectra were measured with a resolution of 60,000, an AGC target of 50% and a mass range from 300 to 1800 m/z. Up to 20 MS2 spectra per duty cycle were triggered at a resolution of 45,000, an AGC target of 300%, an isolation window of 0.7 m/z and a normalized collision energy of 34. Peptides that triggered MS2 scans were dynamically excluded from further MS2 scans for 20s.

### CompRef PDX data analysis

Mass spectrometry data was processed using Spectrum Mill v 7.08 (proteomics.broadinstitute.org). For all samples, extraction of raw files retained spectra within a precursor mass range of 800-6000 Da and a minimum MS1 signal-to-noise ratio of 25. MS1 spectra within a retention time range of +/- 45 s, or within a precursor m/z tolerance of +/- 1.4 m/z were merged. MS/MS searching of PDX samples was performed against a human and mouse RefSeq database with a release date of June 29, 2018 and containing 72,908 entries. Digestion parameters were set to “trypsin allow P” with an allowance of 4 missed cleavages. The MS/MS search included fixed modification of carbamidomethylation on cysteine. For TMT quantitation experiments TMTpro16 was searched using the full-mix function. Variable modifications were acetylation of the protein N-terminus, oxidation of methionine, cyclization to pyroglutamic acid, deamidation, pyrocarbamidomethylation of cysteine and hydroxylation of proline. For PTM datasets, hydroxylation of proline was removed as a variable modification, and additional variable modifications were searched: phosphorylation of serine, threonine and tyrosine residues for IMAC enriched samples; diglycine modification of lysine residues for K(GG) enriched samples. Restrictions for matching included a minimum matched peak intensity of 30% and a precursor and product mass tolerance of +/- 20 ppm.

Peptide spectrum matches were validated using a maximum false discovery rate (FDR) threshold of 0.8% for precursor charges 2 through 4 within each LC-MS/MS run, and 0.4% for precursor charges 5 and 6 within each directory of runs. TMTpro16 reporter ion intensities were corrected for isotopic impurities in the Spectrum Mill protein/peptide summary module using the afRICA correction method which implements determinant calculations according to Cramer’s Rule. We required 2 or more fully quantified unique human peptides with a ratio count of 2 or more for protein identification and a ratio count of 2 or more for protein quantification. To assign regulated proteins and PTM-sites we used the Proteomics Toolset for Integrative Data Analysis (Protigy, v0.9.1.3, Broad Institute, https://github.com/broadinstitute/protigy) to calculate moderated *t*-test *P* values.

### Serial Immunoprecipitation of HLA-I & HLA-II complexes from tumors

Half of each of the ten cryo-pulverized LUAD patient tumors went through the HLA serial immunoprecipitation prior to multi-omic analysis. Each tumor was lysed with 4°C lysis buffer (20mM Tris, pH 8.0, 100mM NaCl, 6mM MgCl2, 1mM EDTA, 60mM Octyl β-d-glucopyranoside, 0.2mM Iodoacetamide, 1.5% Triton X-100, 1xComplete Protease Inhibitor Tablet-EDTA free, 1mM PMSF, 10mM NaF, 1:100 dilution of Protease Inhibitor Cocktail 2 (Sigma-Aldrich, P5726), 1:100 dilution of Protease Inhibitor Cocktail 3 (Sigma-Aldrich, P0044), 50 μMPR-619, 10mM Sodium Butyrate, 2 μMSAHA, 10mM Nicotinamide) obtaining a total of 1.2 ml lysate per tumor. Each lysate was moved into an Eppendorf tube, incubated on ice for 30 min with 2 µL of Benzonase (Thomas Scientific, E1014-25KU) and inverted after 15 min to degrade nucleic acid. The lysates were then centrifuged at 15,000 rcf for 20 min at 4°C and the supernatants were transferred to another set of Eppendorf tubes containing ∼37.5 µL pre-washed Gammabind Plus Sepharose beads(Millipore Sigma, GE17-0886-01). The beads and lysate were rotated at 4°C for one hour in order to preclear hydrophobic molecules and non-specifics that may interfere with the HLA-IP.

The bead-lysate mixtures were centrifuged at 1500 rcf for 1 min at 4°C and each lysate was transferred to a tube containing ∼37.5 µL pre-washed beads and 15 µL of HLA-II antibody mix (9 µL TAL-1B5 (Abcam, ab20181), 3 µL EPR11226 (Abcam, ab157210), 3 µL B-K27 (Abcam, ab47342). The HLA complexes were captured on the beads by incubating on a rotor at 4°C for 3hr. Following the incubation all tubes were centrifuged at 1,500 rcf for 1 min at 4°C and the lysates were transferred from to new Eppendorf tubes containing ∼37.5 µL pre-washed beads and 15 µL of HLA-I antibody (W6/32) (Santa Cruz Biotechnology, sc-32235). The HLA-I antibody/bead/lysate mixture rotated for 3hr at 4°C and was spun at 1500 rcf for 1 min at 4°C. The unbound lysates were transferred to new Eppendorf tubes and flash-frozen with liquid nitrogen for multi-omic downstream analysis.

During HLA complex capture a 10μm PE fritted plate (Agilent, S7898A) was cut in half, placed on a Waters Positive Pressure Manifold, and washed using 1mL acetonitrile and 3x 1mL room-temperature PBS. After each liquid addition, positive pressure of less than 5 psi was applied to the plate to achieve liquid movement. Immediately following each HLA capture, beads were resuspended in 1mL cold PBS and transferred to one half of the pre-washed 10μm PE fritted plate. Each tube was then rinsed with 500 µL cold PBS and remaining beads were transferred to the correct well. In total, four wash steps were performed to remove nonspecifically bound material: two washes with 2mL of cold complete wash buffer (20mM Tris, pH 8.0, 100mM NaCl, 1mM EDTA, 6mM Octyl β-d-glucopyranoside, 0.2mM Iodoacetamide), and two washes with 2mL of 10mM Tris pH 8.0 buffer. The 10μm PE fritted plate with dry HLA-II beads was wrapped with parafilm and stored at 4°C until all HLA-I beads were washed on the other half of the plate and all samples were simultaneously prepared for mass spectrometry analysis via desalting.

### 96-Well Plate Desalt of HLA Immunoprecipitation using a Positive Pressure Manifold

HLA peptides were eluted and desalted from beads as follows: 20 wells of the tC18 40mg Sep-Pak desalting plate (Waters, Milford, MA) were activated two times with 1mL of methanol (MeOH) and 500 µL of 99.9% acetonitrile (ACN)/0.1% formic acid (FA), then washed four times with 1mL of 1% FA. The two halves of the 10μmPE fritted filter plate containing the beads were put together and placed on top of the Sep-Pak plate. To dissociate peptides from HLA molecules and facilitate peptides binding to the tC18 solid-phase, 200 µL of 3% ACN/5% FA was added to the beads in the filter plate. 100fmol internal retention time (iRT) standards (Biognosys SKU: Ki-3002-2) was spiked into each sample as a loading control and pushed through both the filter plate and 40mg Sep-Pak plate. Following sample loading there was one wash with 400 µL of 1% FA. Beads were then incubated with 500 µL of 10% acetic acid (AcOH) three times for 5 min to further dissociate bound peptides from the HLA molecules. The beads were rinsed once with 1mL 1% FA and the filter plate was removed. The Sep-Pak desalt plate was rinsed with 1mL 1% FA an additional three times. The peptides were eluted from the Sep-Pak desalt plate using 250 µL of 15% ACN/1% FA and 2x 250 µL of 50% ACN/1% FA. HLA peptides were eluted into 1.5mL micro tubes (Sarstedt, Nümbrecht, Germany), frozen, and dried down via vacuum centrifugation. Dried peptides were stored at -80°C until microscaled basic reverse phase separation.

Briefly, peptides were loaded on Stage-tips with 2 punches of SDB-XC material (Empore 3M). HLA-I and HLA-II peptides were eluted in three fractions with increasing concentrations of ACN (HLA-I: 5%, 10% and 30% in 0.1% NH4OH, pH 10, HLA-II: 5%, 15%, and 40% in 0.1% NH4OH, pH 10)^23^. Peptides were reconstituted in 3% ACN/5% FA prior to loading onto an analytical column (35 cm, 1.9µm C18 (Dr. Maisch HPLC GmbH), packed in-house PicoFrit 75 µm inner diameter, 10 µm emitter (New Objective)). Peptides were eluted with a linear gradient (EasyNanoLC 1200, Thermo Fisher Scientific) ranging from 6-30% Solvent B (0.1%FA in 90% ACN) over 84 min, 30-90% B over 9 min and held at 90% B for 5 min at 200 nl/min. MS/MS were acquired on a Thermo Orbitrap Exploris 480 equipped with (HLA-I) and without (HLA-II) FAIMS (Thermo Fisher Scientific) in data dependent acquisition. FAIMS CVs were set to -50 and -70 with a cycle time of 1.5s per FAIMS experiment. MS2 fill time was set to 100ms, collision energy was 30CE for HLA-I and 34CE for HLA-II.

### Serial Processing of Ubiquitylome, Proteome, Phosphoproteome, and Acetylome from cryopulverized tissue vs HLA IP flow-through (LUAD)

#### Lysis and tryptic digestion of LUAD cryopulverized tissue and HLA IP flow-through

Each set of 10 replicate tumors underwent denaturing lysis in SDS to prepare for S-Trap digestion. Flow-through of the HLA-I IP, at this point in native HLA lysis buffer and stored as flash-frozen unbound lysates, were briefly thawed on ice for ∼15 min. Once thawed, 10% SDS was added for a final concentration of 2.5% SDS to denature the lysate, resulting in a final volume of ∼1.5 mL lysate which was prepared for S-Trap digestion.

Replicates of the HLA-depleted samples were lysed from cryopulverized tissue in 1 mL 5% SDS buffer (5% SDS, 50 mM TEAB pH 8.5, 2 mM MgCl_2_). The samples were disrupted by pipette mixing and gentle vortexing and incubated at room temperature for ∼10 min. Samples were treated with 2 uL benzonase to shear DNA, mixed again, and incubated at room temperature for another ∼20 min. Finally, non-HLA-depleted lysates were homogenized with a probe sonicator for 30 sec and left to lyse again for ∼10 min. The lysates were cleared by centrifugation for 15 min at 15000 × g and the supernatant was prepared for S-Trap digestion.

In both sets of LUAD tumors, all further processing steps were executed identically. Protein concentration was estimated using a BCA assay for scaling of digestion enzymes. Disulfide bonds were reduced in 5 mM DTT for 30 min at 25°C and 1000 rpm shaking, and cysteine residues alkylated in 10 mM IAA in the dark for 45 min at 25°C and 1000 rpm shaking. Lysates were then transferred to a 15 mL conical tube to prepare for protein precipitation. 27% phosphoric acid was added at a 1:10 ratio of lysate volume to acidify, and proteins were precipitated with 6X sample volume of ice cold S-Trap buffer (90% methanol, 100 mM TEAB). The precipitate was transferred in successive loads of 3 mL to a S-Trap Midi (Protifi) and loaded with 1 min centrifugation at 4000 × g, mixing the remaining precipitate thoroughly between transfers. The precipitated proteins were washed 4x with 3 mL S-Trap buffer at 4000 × g for 1 min. To digest the deposited protein material, 350 µL digestion buffer (50 mM TEAB) containing both trypsin and endopeptidase C (LysC), each at 1:50 enzyme:substrate, was passed through each S-Trap column with 1 min centrifugation at 4000 x g. The digestion buffer was then added back atop the S-Trap and the cartridges were left capped overnight at 25°C.

Peptide digests were eluted from the S-Trap, first with 500 µL 50 mM TEAB and next with 500 µL 0.1% FA, each for 30 sec at 1000 x g. The final elution of 500 µL 50% ACN / 0.1% FA was centrifuged for 1 min at 4000 × g to clear the cartridge. Peptide concentration of the pooled elutions was estimated with a BCA assay, divided into 750 µg peptide aliquots for K-ε-GG enrichment, snap frozen and dried in a vacuum centrifuge.

### Automated UbiFast K-ε-GG enrichment

Peptides containing the K-ε-GG tryptic remnant of ubiquitin/ubiquitin-like small protein modifications were enriched using an adaptation of the UbiFast protocol for the Thermo KingFisher automation platform^20^. Briefly, 750 µg peptide aliquots were reconstituted in 250 µL CST HS bind buffer w/ 0.01% CHAPS. All following steps for UbiFast enrichment excluding labeling and final bead collection contained 0.01% CHAPS. Reconstituted peptides were added to 5 µL PBS-washed CST HS anti-K-ε-GG antibody beads and incubated at 4°C for 1 hour in a foil sealed KingFisher plate with rotation. Following removal of the beads from the incubation by the KingFisher robot, the incubation plate containing non-TMT labeled, K-ε-GG-depleted peptide flow-through was sealed and frozen for downstream proteome, phosphoproteome, and acetylproteome processing. Briefly, bead-bound enriched peptides were washed with 50% CAN / 50% CST HS wash buffer and washed again with PBS. K-ε-GG peptides were labeled on-bead with 400 µg TMT11 in 100 mM HEPES (prepared immediately before run) for 20 minutes and labeling was quenched with 2% hydroxylamine. Finally, the beads were washed with a CST HS wash buffer before being deposited into 100 µL PBS containing no CHAPS buffer. Each well containing each TMT channel was combined by 11-plex, the supernatant was removed, and enriched peptides were eluted from the beads with 2 × 10 min 0.15% TFA. The eluate was desalted with a C18 stagetip, frozen, and dried in a vacuum centrifuge. For LC-MS/MS analysis, the unfractionated K-ε-GG peptides were reconstituted in 9 µL 3% ACN / 0.1% FA and 4 µL was injected twice back-to-back for each sample.

### TMT labeling of UbiFast flow-through for bRP fractionation and analysis proteome

Non-TMT labeled, K-ε-GG-depleted peptide flow-through of the K-ε-GG IPs were acidified with neat formic acid to a final concentration of 1% and desalted with 100 mg tC18 SepPak cartridges. Eluates were frozen and dried in a vacuum centrifuge. Peptides were reconstituted in 30% ACN / 0.1% FA, peptide concentration was estimated using a BCA assay and peptides were aliquoted for downstream processing and dried again. 300 µg of each sample was reconstituted in 60 µL 50 mM HEPES and labeled with 300 µg TMT10 reagent at a final concentration of 20% ACN for 1 hour at 25°C and 1000 rpm. Each tumor replicate was assigned the same TMT channel in its corresponding TMT 10-plex for an identical experimental design. Labeling reactions were diluted to 2.5 mg/mL with 50 mM HEPES. Complete labeling and balancing of input material were confirmed. TMT labeling was quenched with 3 µL 5% hydroxylamine for 15 minutes and each TMT10-plex was combined, frozen, and dried. Dried, labeled, combined peptides were reconstituted with 3 mL 1% FA and desalted with a 200 mg tC18 SepPak, and the eluate was snap frozen and dried in a vacuum centrifuge.

Offline bRP fractionation was performed as described previously and above^17^. Briefly, peptides were separated over a 96 minute gradient with a flow rate of 1 ml/min. Solvent A was 5 mM ammonium formate, 2% ACN and solvent B was 5 mM ammonium formate, 90% ACN. 96 fractions were concatenated into 24 fractions for proteome analysis. 5% of each of the 24 fractions were transferred into HPLC vials, frozen and dried in a vacuum centrifuge for analysis. The remaining 95% of each fraction were concatenated into 13 fractions for phosphopeptide enrichment. Proteome fractions were reconstituted in 3% ACN / 0.1% FA and 500 ng at 0.25 µg/µL from each of the 24 fractions were injected for LC-MS/MS analysis.

### Automated IMAC phosphopeptide enrichment

IMAC enrichment of phosphopeptides was performed using AssayMap Fe(III)-NTA cartridges (Agilent, G5496-60085). Concatenated fractions were solubilized with 80 µL 50% ACN/0.1% TFA in a bath sonicator for 5 min, followed by addition of 120 µL 100% ACN/0.1% TFA for a final concentration of 80% ACN/0.1% TFA. Peptide solution was clarified by centrifugation at 6000 x g for 5 min and 160 µL was transferred to a 96 well plate for enrichment. The remaining 40 µL were set aside for re-enrichment. The Agilent “AssayMAP Phosphopeptide Enrichment v2.1” protocol was used. Briefly, the syringes were rinsed with HPLC water and primed with 50% ACN/0.1% TFA. Cartridges were equilibrated with 80% ACN/0.1% TFA.160 µL of sample was loaded at 5 uL/min, and the phosphopeptide-depleted flow-through was collected and frozen for downstream acetyllysine enrichment. The cartridges were washed 3x with 80% ACN/0.1% TFA to remove non-specific peptides. Enriched phosphopeptides were eluted from the cartridges with 20 µL fresh 1% ammonium hydroxide at 5 uL/min into a plate containing 2.5 µL neat FA. Phosphopeptide-enriched eluates were transferred to HPLC vials, frozen and dried in a vacuum centrifuge. For LC/MS-MS analysis, peptides were reconstituted in 9 µL 3% ACN/0.1% FA and 4 µL were injected from each of the 12 fractions.

### Acetyl-lysine immunoaffinity enrichment

Acetyl peptide enrichment was performed in concordance with^12^ with minor variations described below. Acetylated lysine peptides were enriched using protein A agarose beads conjugated to an acetyl-lysine motif antibody(CST PTM-SCAN Catalog No. 13416). Phosphopeptide-depleted IMAC flow-through were concatenated from 12 to 4 fractions (∼750 μg peptides per fraction) and dried down using a SpeedVac apparatus. Prior to enrichment, antibody beads were washed 4x with IAP buffer (5 mM MOPS pH 7.2, 1 mM sodium phosphate [dibasic], 5 mM NaCl). Peptides were reconstituted with 1.4 mL of IAP buffer per fraction, added to washed beads, and incubated for 2 h at 4°C. Bead-bound acetyl-enriched peptides were washed 4 times with ice-cold PBS followed by two elutions with 100 µLl of 0.15% TFA. Eluents were desalted using C18 stage tips, eluted with 50% ACN / 0.1% FA, and dried down. Acetylpeptides were reconstituted in 7 µL of 3% ACN / 0.1% FA and 4 µL were injected from each of the 4 fractions for LC-MS/MS analysis.

### LC-MS/MS data acquisition of the LUAD samples processed using MONTE

Online separation was done with a nanoflow Proxeon EASY-nLC 1200 UHPLC system (Thermo Fisher Scientific). In this set up, the LC system, column, and platinum wire used to deliver electrospray source voltage were connected via a stainless steel cross (360 mm, IDEX Health & Science, UH-906x). The column was heated to 50°C using a column heater sleeve (Phoenix-ST). Each sample was injected onto an in-house packed 27 cm x 75um internal diameter C18 silica picofrit capillary column (1.9 mm ReproSil-Pur C18-AQ beads, Dr. Maisch GmbH, r119.aq; Picofrit 10 um tip opening, New Objective, PF360-75-10-N-5). Mobile phase flow rate was 200 nL/min, comprised of 3% acetonitrile/0.1% formic acid (Solvent A) and 90% acetonitrile/0.1% formic acid (Solvent B). The same LC and column setup were used for ubiquitylome, proteome, phosphoproteome, and acetylproteome analyses. Each LC-MS/MS method consisted of a 10 min column-equilibration procedure, a 20 min sample-loading procedure, and the following gradient profiles (min:%B): ubiquitylome (154 min) = 0:2, 2:6, 122:35, 130:60, 133:90, 143:90, 144:50, 154:50; proteome/phospho (110 min) = 0:2, 1:6, 85:30, 94:60, 95:90, 100:90, 101:50, 110:50; acetylome (260 min) = 0:2, 1:6, 235:30, 244:60, 245:90, 250:90, 251:50, 260:50. The flow rate of the last two steps of each gradient was increased to 500 nL/min.

For ubiquitylome, proteome, phosphoproteome, and acetylproteome analysis, samples were analyzed with a benchtop Orbitrap Exploris 480 mass spectrometer (Thermo Fisher Scientific) equipped with a NanoSpray Flex NG ion source. Data-dependent acquisition was performed using Orbitrap Exploris 480 V2.0 software in positive ion mode at a spray voltage of 1.8 kV. MS1 spectra were measured with a resolution of 60,000, a normalized AGC target of 300% for proteome/phospho and 100% for ubiquitylome/acetylome, a maximum injection time of 10 ms, and a mass range from 350 to 1800 m/z. The data-dependent mode cycle was set to trigger MS/MS on up to the top 20 most abundant precursors per cycle at an MS2 resolution of 45,000, an AGC target of 30% for proteome/phospho and 50% for ubiquitylome/acetylome, an isolation window of 0.7 m/z, a maximum injection time of 105 ms for proteome/phospho and 120 ms for ubiquitylome/acetylome, and an HCD collision energy of 34%. Peptides that triggered MS/MS scans were dynamically excluded from further MS/MS scans for 20 s in proteome/phospho/ubiquitylome and for 30 s in acetylome, with a ±10 ppm mass tolerance. Theoretical precursor envelope fit filter was enabled with a fit threshold of 50% and window of 1.2 m/z. Monoisotopic peak determination was set to peptide, and charge state screening was enabled to only include precursor charge states 2-6, with an intensity threshold of 5.0e3. Advanced peak determination (APD) was enabled. “Perform dependent scan on single charge state per precursor only” was disabled.

### LUAD MONTE LC-MS/MS data interpretation

MS/MS spectra from all ‘omes were interpreted using Spectrum Mill (SM) v 7.08 (proteomics.broadinstitute.org) to provide identification and relative quantitation at the protein, peptide, and post-translational modification (PTM) site (ubiquityl, phospho, and acetyl) site levels.

### Variant Calls

Individual variant/indel .vcf files for each of the 10 LUAD patients in this study were extracted from the CPTAC Pancancer Harmonized Callset v1.1 which is the harmonized result of processing whole exome sequencing data from 10 CPTAC cancer cohorts independently through the variant calling pipelines of the Getz laboratory at the Broad Institute and the Ding lab at Washington University in St Louis. The Getz laboratory pipeline consists of GATK(v4.1.4.1) for DNA sequence data quality control and somatic copy number analysis, MuTect^42^ Manta+Strelka v2^69,70^ for discovery of somatic and germline SNVs and INDELs, DeTiN v1.8.9^71^ and GATK4 Funcotator ver GATK 4.1.4.1 for post-discovery filtering followed by merging of adjacent somatic SNPs into DNPs, TNPs and ONPs. The Ding laboratory employed the Somaticwrapper pipeline v1.6 (https://github.com/ding-lab/somaticwrapper), which includes four different callers, i.e., Strelka v.2^69,72^, MUTECT v1.7^42^, VarScan v.2.3.8^73^, and Pindel v.0.2.574. Rare mutations with VAF of [0.015, 0.05) in cancer driver genes were rescued based on the gene consensus list reported in^75^. COCOON (https://github.com/ding-lab/COCOONS) was used to combine adjacent SNVs into DNPs.

### Personalized sequence database

For searching with LC-MS/MS datasets from all ‘omes we generated a personalized protein sequence database starting with a base human reference proteome to which we appended somatic and germline variants and indels for each of the 10 LUAD patients. The base proteome consisted of the human reference proteome Gencode 34 (ftp.ebi.ac.uk/pub/databases/gencode/Gencode_human/release_34/) with 47,429 non-redundant protein coding transcript biotypes mapped to the human reference genome GRCh38, 602 common laboratory contaminants, 2043 curated smORFs (lncRNA and uORFs), 237,427 novel unannotated ORFs (nuORFs) supported by ribosomal profiling nuORF DB v1.0^37^, and 4,167 TCGA shared mutations from 26 tumor types (https://www.cancer.gov/tcga) for a total of 355,028 entries which yield 16,973,937 distinct 9-mers. The nuORFs alone yield 8,612,372 distinct 9-mers and thus increase the peptide search space by only a factor of ∼2.

The personalized protein sequence entries were prepared by processing the individual patient’s somatic and germline variant calls from whole exome sequencing data, described above, using QUILTS v3^42–44^ with no further variant quality filtering using a Ensembl v100 reference proteome and reference genome for sequence identifiers consistent with the variant calling. Gencode v34 is a contemporaneous subset of Ensembl v100(March 2020). Using the SM Protein Database utilities the base reference proteome and individual patient proteomes were combined and redundancy removed to produce a cohort-level protein sequence database and a variant summary table to enable subsequent mapping of sequence variants identified in TMT multiplexed LC-MS/MS datasets back to individual patients.

### Spectrum quality filtering

Using the SM Data Extractor module for HLA I,II ‘omes spectral merging was disabled, the precursor MH+ inclusion range was 600-4000, and the spectral quality filter was a sequence tag length >1 (i.e., minimum of three peaks separated by the in-chain masses of two consecutive amino acids). For non-HLA ‘omes, similar MS/MS spectra with the same precursor m/z acquired in the same chromatographic peak were merged, the precursor MH+ inclusion range was 800-6000, and the spectral quality filter was a sequence tag length > 0.

### MS/MS search conditions

Using the SM MS/MS search module for HLA I,II ‘omes parameters included: no-enzyme specificity; precursor and product mass tolerance of ±10 ppm; minimum matched peak intensity of 30%; ESI-QEXACTIVE-HCD-HLA-v3 scoring; fixed modification: carbamidomethylation of cysteine; variable modifications: cysteinylation of cysteine, oxidation of methionine, deamidation of asparagine, acetylation of protein N-termini, and pyroglutamic acid at peptide N-terminal glutamine; precursor mass shift range of -18 to 81 Da. For HLA I,II a second round search of remaining unassigned spectra was done with variable modifications revised to also allow for acetylation of lysine and phosphorylation of serine, threonine, and tyrosine with a precursor MH+ shift range of -18 to 125 Da.

For non-HLA ‘omes parameters included: ‘‘trypsin allow P’’ enzyme specificity with up to 4 missed cleavages; precursor and product mass tolerance of ± 20 ppm; 30% minimum matched peak intensity (40% for acetylome). Scoring parameters were ESI-QEXACTIVE-HCD-v2, for whole proteome datasets, and ESI-QEXACTIVE-HCD-v3, for phosphoproteome, acetylome, and ubiquitylome datasets. Allowed fixed modifications included carbamidomethylation of cysteine and selenocysteine. TMT labeling was required at lysine, but peptide N-termini were allowed to be either labeled or unlabeled. Allowed variable modifications for whole proteome datasets were acetylation of protein N-termini, oxidized methionine, deamidation of asparagine, hydroxylation of proline in PG motifs, pyro-glutamic acid at peptide N-terminal glutamine, and pyro-carbamidomethylation at peptide N-terminal cysteine with a precursor MH+ shift range of - 18 to 97 Da. For all PTM-omes variable modifications were revised to omit hydroxylation of proline and allow deamidation only in NG motifs. The phosphoproteome was revised to allow phosphorylation of serine, threonine, and tyrosine with a precursor MH+ shift range of -18 to 272 Da. The acetylome was revised to allow acetylation of lysine with a precursor MH+ shift range of -400 to 70 Da. The ubiquitylome was revised to allow diglycine modification of lysine with a precursor MH+ shift range of -375 to 70 Da.

### PTM site localization

Using the SM Autovalidation and Protein/Peptide Summary modules for the PTM-ome datasets results were filtered and reported at the ubiquityl, phospho, and acetyl site levels. When calculating scores at the variable modification (VM) site level and reporting the identified VM sites, redundancy was addressed in SM as follows: a VM-site table was assembled with columns for individual TMT-plex experiments and rows for individual VM-sites. PSMs were combined into a single row for all non-conflicting observations of a particular VM-site (e.g., different missed cleavage forms, different precursor charges, confident and ambiguous localizations, and different sample-handling modifications). For related peptides, neither observations with a different number of VM-sites nor different confident localizations were allowed to be combined. Selecting the representative peptide for a VM-site from the combined observations was done such that once confident VM-site localization was established, higher identification scores and longer peptide lengths were preferred. While an SM PSM identification score was based on the number of matching peaks, their ion type assignment, and the relative height of unmatched peaks, the VM site localization score was the difference in identification score between the top two localizations. The score threshold for confident localization, > 1.1, essentially corresponded to at least 1 b or y ion located between two candidate sites that has a peak height > 10% of the tallest fragment ion (neutral losses of phosphate from the precursor and related ions as well as immonium and TMT reporter ions were excluded from the relative height calculation). The ion type scores for b-H3PO4, y-H3PO4, b-H2O, and y-H2O ion types were all set to 0.5. This prevented inappropriate confident localization assignment when a spectrum lacked primary b or y ions between two possible sites but contained ions that could be assigned as either phosphate-loss ions for one localization or water loss ions for another localization.

### Protein grouping of PSMs, peptides and PTM sites

Using the SM Autovalidation and Protein/Peptide summary modules results were filtered and reported at the protein level. Identified proteins were combined into the same protein group if they shared a peptide with sequence length greater than 8. A protein group could be expanded into subgroups (isoforms or family members) when distinct peptides were present which uniquely represent a subset of the proteins in a group. For the proteome dataset the protein grouping method ‘‘expand subgroups, top uses shared’’ (SGT) was employed which allocates peptides shared by protein subgroups only to the highest scoring subgroup containing the peptide. For the PTM-ome datasets the protein grouping method ‘‘unexpand subgroups’’ was employed which reports a VM-site only once per protein group allocated to the highest scoring subgroup containing the representative peptide. The SM protein score is the sum of the scores of distinct peptides. A distinct peptide is the single highest scoring instance of a peptide detected through an MS/MS spectrum. MS/MS spectra for a particular peptide may have been recorded multiple times (e.g., as different precursor charge states, in adjacent bRP fractions, modified by deamidation at Asn or oxidation of Met, or with different phosphosite localization), but are still counted as a single distinct peptide.

### Peptide spectrum match (PSM) filtering and false discovery rates (FDR)

Using the SM Autovalidation module peptide spectrum matches (PSMs) for individual spectra were confidently assigned by applying target-decoy based FDR estimation to achieve <1.0% FDR at the PSM, peptide, VM site and protein levels. For HLA I,II ‘omes PSM level thresholding was done with a minimum peptide length of 7, minimum backbone cleavage score of 5, and <1.0% FDR across all 3 fractions. Allowed precursor charges were HLA-I : 1-4, HLA-II: 2-6. Immunopeptidomics data was further filtered to remove non-human contaminants, peptides that match peptides identified in blank bead negative control IPs^7,8^, and tryptic contaminant peptides. Phospho- and acetyl HLA peptides were quality filtered to include matches with scores >6 and scored peak intensity >60%; HLA-I included only 8-11mers.

For the whole proteome dataset thresholding was done in 3 steps: at the PSM level, the protein level for each TMT-plex, and the protein level for the cohort of 2 TMT-plexes obtained with and without initial HLA IP. For the PTM-omes: ubiquitylome, phosphoproteome, and acetylome datasets thresholding was done in two steps: at the PSM level for each TMT-plex and at the VM site level for the cohort of 2 TMT-plexes. In step 1 for all datasets, PSM level autovalidation was done first and separately for each TMT-plex experiment using an auto-thresholds strategy with a minimum sequence length of 7; automatic variable range precursor mass filtering; with score and delta Rank1 - Rank2 score thresholds optimized to yield a PSM level FDR estimate for precursor charges 2 through 4 of < 0.8% for each precursor charge state in each LC-MS/MS run. To achieve reasonable statistics for precursor charges 5-6, thresholds were optimized to yield a PSM-level FDR estimate of < 0.4% across all runs per TMT-plex experiment (instead of per each run), since many fewer spectra are generated for the higher charge states.

In step 2 for the PTM-omes: ubiquitylome, phosphoproteome, and acetylome datasets VM site polishing autovalidation was applied across both TMT plexes to retain all VM site identifications with either a minimum id score of 8.0 or observation in both TMT plexes. The intention of the VM site polishing step is to control FDR by eliminating unreliable VM site level identifications, particularly low scoring VM sites that are only detected as low scoring peptides that are also infrequently detected across both TMT plexes in the study. Using the SM Protein/Peptide Summary module to make VM-site reports the ubiqiuitylome and acetylome datasets were further filtered to remove peptides ending with the regular expression [^K][^K]k since trypsin and Lys-C cannot cleave at a ubiquitylated or acetylated lysine. The [^K] means retain if unmodified Lys present in one of the last two positions to allow for a missed cleavage with ambiguous PTM-site localization.

In step 2 for the whole proteome dataset, protein polishing autovalidation was applied separately to each TMT-plex experiment to further filter the PSMs using a target protein level FDR threshold of zero. The primary goal of this step was to eliminate peptides identified with low scoring PSMs that represent proteins identified by a single peptide, so-called ‘‘one-hit wonders.’’ After assembling protein groups from the autovalidated PSMs, protein polishing determined the maximum protein level score of a protein group that consisted entirely of distinct peptides estimated to be false-positive identifications (PSMs with negative delta forward-reverse scores). PSMs were removed from the set obtained in the initial peptide level autovalidation step if they contributed to protein groups that had protein scores below the maximum false-positive protein score. Step 3 was then applied, consisting of protein polishing autovalidation across both TMT plexes together using the protein grouping method ‘‘expand subgroups, top uses shared’’ to retain protein subgroups with either a minimum protein score of 25 or observation in both TMT plexes. The primary goal of this step was to eliminate low scoring proteins that were infrequently detected in the sample cohort. As a consequence of these two protein-polishing steps, each identified protein reported in the study comprised multiple peptides, unless a single excellent scoring peptide was the sole match and that peptide was observed in both TMT-plexes.

### Subset-specific FDR filtering for neoantigens, nuORFs and somatic variants

All MS/MS spectra of neoantigens were manually inspected and labeled spectra are provided in **Figure 5D** and **Figure S2**. While the aggregate FDR for each dataset was set to < 1%, as described above, FDR for certain subsets of rarely observed classes (<5% of total) of peptides, PTM sites, and proteins required more stringent score thresholding to reach a suitable subset-specific FDR < 1.0%. To this end, we devised and applied subset-specific filtering approaches.

Subsets of nuORF types were thresholded independently in the HLA and PTM-ome datasets using a 2-step approach. First, PSM scoring metric thresholds were tightened in a fixed manner for all nuORF PSMs so that nuORF distributions for each metric improved to meet or exceed the aggregate distributions. For all ‘omes the fixed thresholds were: minimum score: 7, minimum percent scored peak intensity: 50%, precursor mass error: +/-5 ppm. For HLA ‘omes minimum backbone cleavage score (BCS): 5, sequence length: 8-12 (HLA-I), 9-50 (HLA-II). For PTM-omes these fixed thresholds were: minimum score: 7, minimum backbone cleavage score (BCS): 4, sequence length: 7-50. Second, individual nuORF type subsets with FDR estimates remaining above 1% were further subject to a grid search to determine the lowest values of BCS (sequence coverage metric) and score (fragment ion assignment metric) that improved FDR to < 1% for each ORF type in the dataset for each ome.

The subset of peptides containing single amino acid variants (SAAVs) and indels observed in the proteome was extracted after step 1 of PSM filtering described above using the SM Protein/Peptide Summary module to create a proteogenomics (PG) site report, with quantitation normalized to nullify the effect of differential protein loading using the aggregate protein-level normalization factors from the fully filtered proteome dataset. The PG-site report was manually filtered to the final subset of somatic SAAVs and indels by retaining those in which the TMT ratios were extremely high only for the patients in which the corresponding SNV or indel was observed.

### Quantitation using TMT ratios

Using the SM Protein/Peptide Summary module, a protein comparison report was generated for the proteome dataset using the protein grouping method ‘‘expand subgroups, top uses shared’’ (SGT). For the PTM-omes: ubiquitylome, phosphoproteome, and acetylome datasets VM site comparison reports limited to either ubiquityl, phospho, or acetyl sites, respectively, was generated using the protein grouping method ‘‘unexpand subgroups.’’ Relative abundances of proteins and VM-sites were determined in SM using TMT reporter ion log2 intensity ratios from each PSM. TMT reporter ion intensities were corrected for isotopic impurities in the SM Protein/Peptide Summary module using the afRICA correction method, which implements determinant calculations according to Cramer’s Rule and correction factors obtained from the reagent manufacturer’s certificate of analysis (https://www.thermofisher.com/order/catalog/product/90406) for TMT10 lot number UA280170. Each protein-level or PTM site-level TMT ratio was calculated as the median of all PSM-level ratios contributing to a protein subgroup or PTM site. PSMs were excluded from the calculation if they lacked a TMT label, had a precursor ion purity < 50% (MS/MS has significant precursor isolation contamination from co-eluting peptides), or had a negative delta forward-reverse identification score (half of all false-positive identifications). Using the SM Process Report module non-quantifiable proteins and PTM sites (ex: unlabeled peptides containing an acetylated protein N-terminus and ending in arginine rather than lysine) were removed, and median/MAD normalization was performed on each TMT channel in each ome to center and scale the aggregate distribution of protein-level or PTM site-level log-ratios around zero in order to nullify the effect of differential protein loading and/or systematic MS variation. When subsets of an ome (nuORF or SAAVs, etc) the TMT ratios were normalized using the normalization factors for the aggregate distribution of the corresponding ome.

### HLA peptide prediction using HLAthena

HLA peptide prediction was performed using HLAthena ^8^. Unless otherwise specified, peptides were assigned to an allele using a percentile rank cutoff ≤ 0.5.

## Data Availability

The original mass spectra and the protein sequence databases used for searches have been deposited in the public proteomics repository MassIVE (http://massive.ucsd.edu). These datasets will be made public upon acceptance of the manuscript.

## Acknowledgements

We thank Shankha Satpathy for thoughtful discussions related to experimental design and analysis, and Cadence Pearce for contributions related to preliminary studies not shown in this manuscript. We thank Yo Akiyama, Qing Zhang, Francois Aguet, Yifat Geffen, and Matthew Wyczalkowski for performing somatic and germline variant calling. This work was supported by the National Cancer Institute (NCI) grants U24CA210986, U01CA214125, and U24CA210979 to S.A.C., Swiss National Science Foundation (SNF) Sinergia grant CRSII5_186405 to S.A.C., Dr. Miriam and Sheldon G. Adelson Medical Research Foundation to S.A.C. and N.D.U. and by a SPARC Award to N.D.U. from the Broad Institute of MIT & Harvard (#800373). This work was also made possible in part by a grant from BroadIgnite at the Broad Institute of MIT and Harvard to S.K.

