## Supplemental for "MONTE enables serial immunopeptidome, ubiquitylome, proteome, phosphoproteome, acetylome analyses of sample-limited tissues"

A.

| sample_annotation | DPA1-1 | DPA1-2 | DPB1-1 | DPB1-2 | DQA1-1 | DQA1-2 | DQB1-1 | DQB1-2 | DRA1-1 | DRA1-2 | DRB1-1 | DRB1-2 | DRB3-1 | DRB3-2 | DRB4-1 | DRB4-2 | DRB5-1 | DRB5-2 |
|---|---|---|---|---|---|---|---|---|---|---|---|---|---|---|---|---|---|---|
| C3L-01632 | DPA1*01:03:01 | DPA1*01:03:01 | DPB1*02:01:02 | DPB1*04:01:01 | DQA1*05:05:01 | DQA1*01:02:01 | DQB1*06:04:01 | DQB1*03:01:01 | DRA*01:01:01 | DRA*01:02:02 | DRB1*11:01:01 | DRB1*13:02:01 | DRB3*03:01:01 | DRB3*02:02:01 | NA | NA | NA | NA |
| C3L-02549 | DPA1*01:03:01 | DPA1*01:03:01 | DPB1*15:01:01 | DPB1*04:01:01 | DQA1*01:03:01 | DQA1*01:02:01 | DQB1*06:01:01 | DQB1*06:02:01 | DRA*01:02:02 | DRA*01:02:03 | DRB1*15:02:01 | DRB1*15:01:01 | NA | NA | NA | NA | DRB5*01:01:01 | DRB5*01:02:01 |
| C3N-00169 | DPA1*04:01:01 | DPA1*02:02:02 | DPB1*107:01 | DPB1*05:01:01 | DQA1*03:02:01 | DQA1*01:02:02 | DQB1*05:02:01 | DQB1*03:03:02 | DRA*01:01:01 | DRA*01:01:01 | DRB1*16:02:01 | DRB1*09:01:02 | NA | NA | NA | DRB4*01:03 | NA | DRB5*01:01:01 |
| C3N-00199 | DPA1*02:02:02 | DPA1*01:03:01 | DPB1*01:01:01 | DPB1*04:01:01 | DQA1*02:01:01 | DQA1*01:02:02 | DQB1*02:02:01 | DQB1*05:02:01 | DRA*01:01:01 | DRA*01:01:01 | DRB1*16:01:01 | DRB1*07:01:01 | NA | NA | NA | DRB4*01:03 | NA | DRB5*02:02:01 |
| C3N-00547 | DPA1*02:02:02 | DPA1*01:03:01 | DPB1*05:01:01 | DPB1*02:01:02 | DQA1*06:01:01 | DQA1*03:01:01 | DQB1*03:02:01 | DQB1*03:01:01 | DRA*01:01:01 | DRA*01:02:02 | DRB1*12:02:01 | DRB1*04:04:01 | NA | DRB3*03:01:03 | NA | DRB4*01:03 | NA | NA |
| C3N-00579 | DPA1*04:01:01 | DPA1*01:03:01 | DPB1*107:01 | DPB1*03:01:01 | DQA1*06:01:01 | DQA1*03:02:01 | DQB1*03:01:01 | DQB1*03:03:02 | DRA*01:01:01 | DRA*01:02:02 | DRB1*12:02:01 | DRB1*09:01:02 | NA | DRB3*03:01:03 | NA | DRB4*01:03 | NA | NA |
| C3N-01016 | DPA1*01:03:01 | DPA1*01:03:01 | DPB1*04:01:01 | DPB1*04:01:01 | DQA1*05:01:01 | DQA1*03:03:01 | DQB1*04:01:01 | DQB1*02:01:01 | DRA*01:01:01 | DRA*01:01:01 | DRB1*04:05:01 | DRB1*03:01:01 | DRB3*02:02:01 | NA | DRB4*01:03 | NA | NA | NA |
| C3N-01024 | DPA1*02:02:02 | DPA1*02:02:02 | DPB1*05:01:01 | DPB1*05:01:01 | DQA1*06:01:01 | DQA1*06:01:01 | DQB1*03:116 | DQB1*03:01:01 | DRA*01:02:02 | DRA*01:02:02 | DRB1*12:02:01 | DRB1*12:02:01 | DRB3*03:01:03 | DRB3*03:01:03 | NA | NA | NA | NA |
| C3N-01416 | DPA1*04:01:01 | DPA1*02:01:01 | DPB1*107:01 | DPB1*13:01:01 | DQA1*06:01:01 | DQA1*03:02:01 | DQB1*03:03:02 | DQB1*03:01:01 | DRA*01:01:01 | DRA*01:02:02 | DRB1*12:02:01 | DRB1*09:01:02 | NA | DRB3*03:01:03 | NA | DRB4*01:03 | NA | NA |
| C3N-02145 | DPA1*02:02:02 | DPA1*02:02:02 | DPB1*05:01:01 | DPB1*05:01:01 | DQA1*06:01:01 | DQA1*05:03:01 | DQB1*03:01:01 | DQB1*03:116 | DRA*01:01:01 | DRA*01:02:02 | DRB1*12:02:01 | DRB1*13:12:01 | DRB3*03:01:03 | DRB3*02:02:01 | NA | NA | NA | NA |

B.

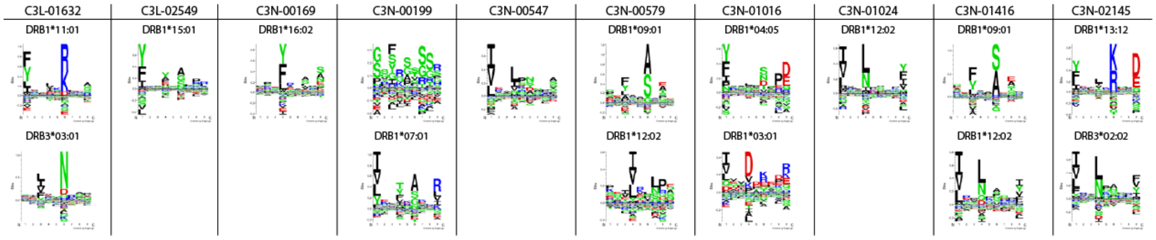


**Figure S1: HLA-II alleles and corresponding peptide binding motifs identified from LUAD immunopeptidomes. A**)HLA-II alleles expressed by the LUAD patient cohort that were called from RNAseq data using arcasHLA[4]. HLA-DRB4 alleles (red) were imputed from known genomic linkages[5,6] because no calls were made for these alleles by arcasHLA. **B**)HLA-II binding motifs that were found using Gibbs Cluster[7] and could be assigned to patient HLA-II alleles.

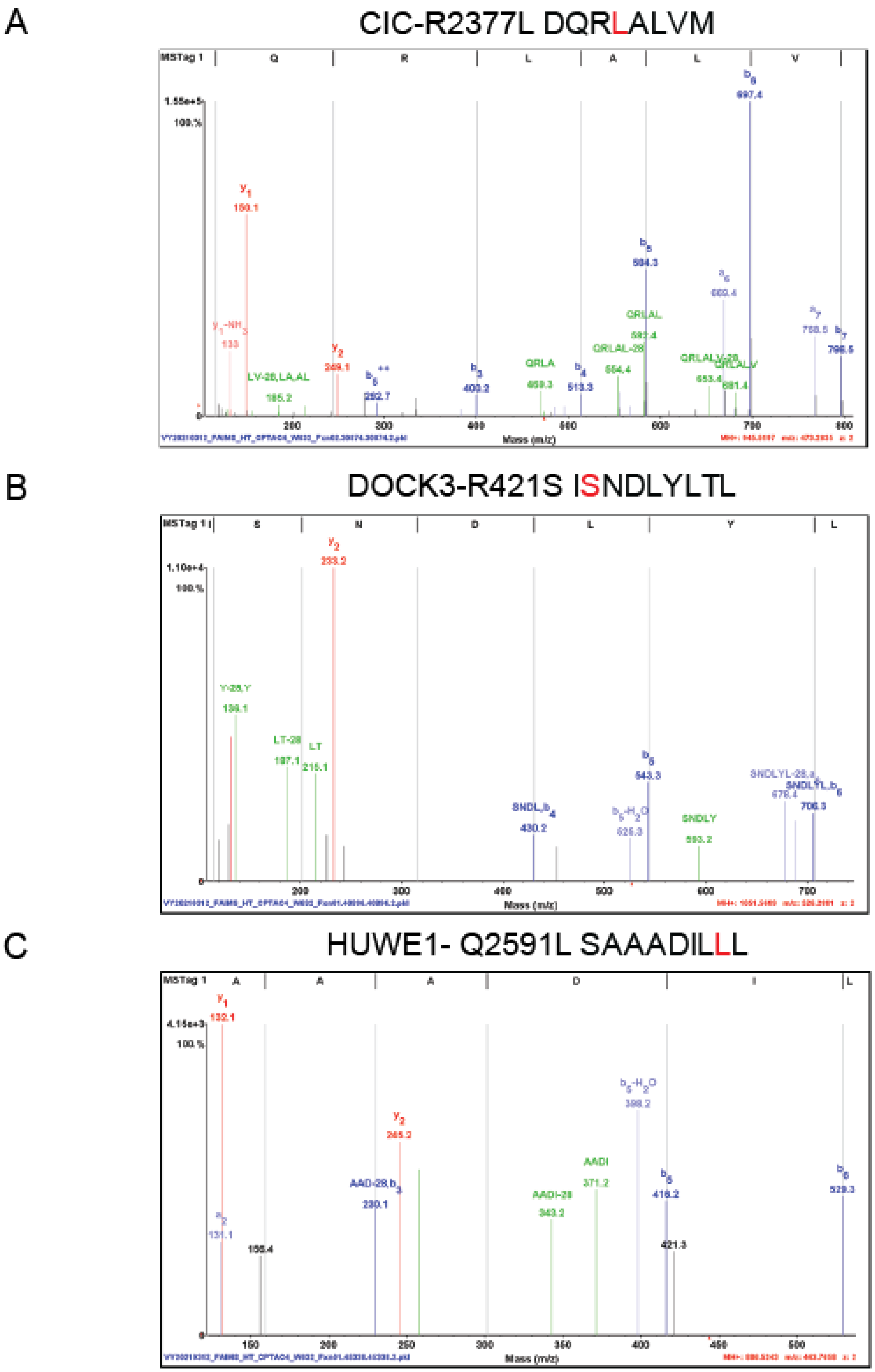


**Figure S2:** Annotated MS/MS spectra for neoantigens detected in LUAD patient C3N-00547. **A**) CIC R2377L neoantigen peptide DQRLALVM. **B**) DOCK3 R421S neoantigen peptide ISNDLYLTL. **C**) HUWE1 Q2591L neoantigen peptide SAAADILLL.
